# Cross-Species BAC Transgenesis Reveals Long-Range Regulation Drives Variation in Brain Oxytocin Receptor Expression and Social Behaviors

**DOI:** 10.1101/2022.12.01.518660

**Authors:** Mina Tsukamoto, Luis AE Nagai, Kiyoshi Inoue, Lenin C. Kandasamy, Maria F. Pires, Minsoo Shin, Yutaro Nagasawa, Tsetsegee Sambuu, Kenta Nakai, Shigeyoshi Itohara, Larry J Young, Qi Zhang

**Author notes:** Deceased in March, 2024. We dedicate this work to the memory of Dr. Larry J. Young, whose encouragement and unwavering support helped our team persevere through challenges.

## Abstract

Although oxytocin (OXT) exhibits a highly conserved neuroanatomical pattern among vertebrates, the distribution of OXT receptor (OXTR) in brain varies considerably across species and is associated with species-typical social behavior. To investigate the genomic basis of the phylogenetic plasticity in brain *Oxtr* expression and its social behavioral consequences, we generated transgenic mice carrying a bacteria artificial chromosome (BAC) harboring the entire prairie vole *Oxtr* locus and flanking intergenic regulatory regions. We established eight independent “volized” mouse lines expressing prairie vole *Oxtr* (pv*Oxtr*). Strikingly, despite conserved *Oxtr* expression in mammary gland of all transgenic mouse lines, each line displayed a unique pattern of brain expression distinct from both mice and prairie voles. Together with topologically associating domain (TAD) structure analysis with mouse genome, our findings suggest that unlike *Oxt*, *Oxtr* expression patterns in brain, involve contributions of distal regulatory elements beyond the BAC insert. In contrast, *Oxtr* expression in peripheral tissues appears resistant to such distal influences. Moreover, the “volized” mouse lines with different brain *Oxtr* expression patterns showed differences in partner preference and maternal behaviors, providing direct functional evidence that variation in brain *Oxtr* expression can drive differences in social behaviors. We propose that brain *Oxtr* expression is transcriptionally sensitive to long-range interactions with distal genomic elements, rendering it more susceptible to diverse regulatory influences. This supports a model in which regulatory flexibility facilitates the evolutionary diversification of social behavior, while maintaining essential peripheral *Oxtr* expression.

## Introduction

Brain oxytocin receptors (OXTR) regulate a wide range of social behaviors including social recognition, maternal care, social bonding, empathy related behaviors, and aggression (Jurek and Neumann, 2018; Froemke and Young, 2021; Rigney et al., 2022). In contrast to sex steroid receptors, which have highly conserved brain expression patterns across species, OXTR shows remarkable inter- and intra-species variation in brain distribution (Froemke and Young, 2021; Young and Zhang, 2021). For example, monogamous prairie voles have a brain OXTR distribution in cortex, striatum and amygdala that is significantly distinct from promiscuous montane voles and laboratory rats and mice(Young and Zhang, 2021). Furthermore, there is robust individual variation in brain OXTR expression among prairie voles that is associated with single nucleotide polymorphisms (SNPs) in the OXTR locus (*Oxtr*), and this variation predicts pair bonding behavior and resilience to early life social neglect (Ross et al., 2009; Barrett et al., 2015; King et al., 2016; Ahern et al., 2021). Different species of primates also differ in brain OXTR distribution (Rogers Flattery et al., 2022), and SNPs in the human OXTR have been linked to variation in social function (Skuse et al., 2014; Theofanopoulou et al., 2022). Thus, variation in OXTR distribution in brain likely constitutes a mechanism for the emergence of diverse social traits, potentially including psychiatric endophenotypes.

The evolution of gene expression patterns could be mediated by *cis* (via linked polymorphisms) or *trans* (through diffusible products of distal genes, e.g., transcription factors) changes, as a result of adaptation to the environment (Shi et al., 2012; Mack et al., 2016; Osada et al., 2017). *Cis*-regulatory differences are more commonly responsible for adaptive evolution and interspecific divergence (Signor and Nuzhdin, 2018). To explore the transcriptional mechanisms giving rise to species-specific *Oxtr* expression and social behavior, we created transgenic mice using a prairie vole *Oxtr* (pv*Oxtr*) bacterial artificial chromosome (BAC). The BAC construct covers the full coding sequence, introns and the entire intergenic region of the pv*Oxtr* with its neighboring genes. We expected that our BAC construct accommodates all the critical regulatory sequences responsible for species- and tissue-specific expression, so that pv*Oxtr* transgenic mice would express brain pv*Oxtr* in a prairie vole-like pattern, largely independent of integration sites. Then we would examine if our transgenic mouse could develop social behaviors similar to that of prairie vole.

Contrary to our initial expectation, the independent Koi lines did not reproduce a single prairie vole-like *Oxtr* expression pattern but instead exhibited distinct and partially overlapping pvOXTR expression configurations. This unexpected diversity provided an opportunity to address a broader question: how does variation in *Oxtr* regulatory architecture generate diversity in receptor expression patterns and, consequently, social behaviors? We therefore used the Koi lines to examine whether divergent pvOXTR expression patterns are associated with differences in social behavior within the same mouse genetic background, rather than to identify a Koi line that fully recapitulates prairie vole social behavior.

We generated eight pv*Oxtr* mouse lines with successful germline transmission. Each line, defined by a distinct BAC integration site, exhibited a unique brain-specific expression pattern of *Oxtr*, distinct from both wildtype mice and prairie voles. Notably, *Oxtr* expression in the mammary gland was conserved across all lines. This suggests that brain, but not mammary gland, expression of *Oxtr* is shaped by distal regulatory elements beyond the BAC insert. These findings imply that evolution has placed brain *Oxtr* expression under more complex regulatory control, likely to accommodate the diversification of social behaviors, whereas mammary gland expression is maintained by simpler mechanisms supporting conserved lactation functions. Furthermore, behavioral assays revealed line-specific differences in partner preference and maternal behaviors.

The 3D architecture of the genome, segregated into TADs demarcated by CCCTC-binding factor (CTCF), crucially influences gene expression. Through the loop extrusion model, CTCF enables interactions between distant regulatory elements, facilitating precise gene regulation, which ultimately shapes an organism’s phenotype and behavior(Wendt et al., 2008). Hi-C data provide a genome-wide view of chromosome conformation in multiple layers, such as compartments, TADs, and loops(Lieberman-Aiden et al., 2009; Gibcus and Dekker, 2013). Our TAD analysis with Hi-C database showed a clear difference exists between the 3D structures of *Oxt* and *Oxtr* gene, and a clear difference exists between the 3D structures of brain tissue *Oxtr* and peripheral tissue *Oxtr*, suggesting that Oxtr expression patterns in brain depends on contributions from very distal sequences. These results provide important clues to understand the genetic mechanism underlying species differences in brain gene expression.

Together, our data demonstrate that transcriptional redistribution of brain *Oxtr* can drive variation in social and parental behaviors—variation that natural selection could act upon in specific ecological contexts. Our results also offer insights into the genetic mechanisms underlying species-specific brain gene expression.

## Results

### Creation of the pv*Oxtr*-*P2A*-*Cre* BAC transgenic mice

BAC vectors can accommodate dispersed cis-regulatory elements across large regions of genome. Transgene expression from BAC constructs are generally resistant to insertion position effects (Suster et al., 2011) due to both the large spans of insulating genetic material which protect the transgene cassette from the influence of the chromosomal environment, and through the inclusion of necessary regulatory elements such as enhancers, silencers, locus control regions, and matrix attachment regions (Bian and Belmont, 2010; Beil et al., 2012). Large-scale screening of CNS gene expression in BAC transgenic mice by the GENSAT (Gene Expression Nervous System Atlas) Project found that more than 85% of BACs express reproducibly in multiple independent transgenic lines and reporter gene expression faithfully recapitulates endogenous gene expression patterns (Heintz, 2004; Gerfen et al., 2013; Schmidt et al., 2013). Indeed, reproducible expression from mouse BAC vectors has been achieved for more than 500 genes (Heintz, 2004; Gerfen et al., 2013; Schmidt et al., 2013). Few that don’t express reproducibly are influenced by variation in BAC copy number and insertion sites (Gong et al., 2007). BAC engineering has also been used to mediate cross-species transgene expression. A large number of mouse models of human dominant neurodegenerative disorders have been created using human BAC transgene approaches. These model mice successfully express human transgenes in human-like patterns and recapitulate disease-like phenotypes (Yang and Lu, 2008; Ferrante, 2009; Crook and Housman, 2011; Taguchi et al., 2019; Shenoy et al., 2022). Previously we constructed *Pkcd-BAC-Cre* transgenic mice and obtained independent lines with reproducible expression patterns as well (Zhang et al., 2016)(Supplementary Fig.1).We therefore decided to use BAC approaches to create “volized” *Oxtr* mouse lines.

To create pv*Oxtr*-*P2A*-*Cre* BAC transgenic mouse line, the prairie vole BAC clone we used includes the entire pv*Oxtr* locus (the coding sequence and all the introns) with the full intergenic sequence upstream (150kb) and downstream of the pv*Oxtr* coding region (9kb) (Fig.1 B). The BAC clone contains a portion of *Rad18,* minus the first 11 exons, located upstream of pv*Oxtr,* and *Cav3*, minus the first exon including the start codon, downstream of pv*Oxtr.* Therefore, there will be no interference of these two flanking transgenes on behavior analysis, and the proximal topological structure around pv*Oxtr* locus is preserved to the maximum extent. A P2A-NLS-Cre cassette was inserted in-frame just before the *Oxtr* stop codon to ensure co-expression of *Cre* and *Oxtr* (Fig.1 C). Transgenic mice were generated by pronuclear injection using C57BL/6J zygotes. Proper integration of the reporter into the BAC clone was confirmed using gene specific PCR assays and sequencing (Fig.1 C, D and data not shown). We obtained 11 founder mice carrying the transgenes, which was verified by PCR with multiple primer sets (Fig.1 C, E). One founder line was sterile, and two founder lines did not show germ-line transmission. Eight founder lines successfully produced offspring and stably transmitted the transgene to at least 3 generations. We named these lines “Koi”, meaning “love” in Japanese.

**Fig. 1.**
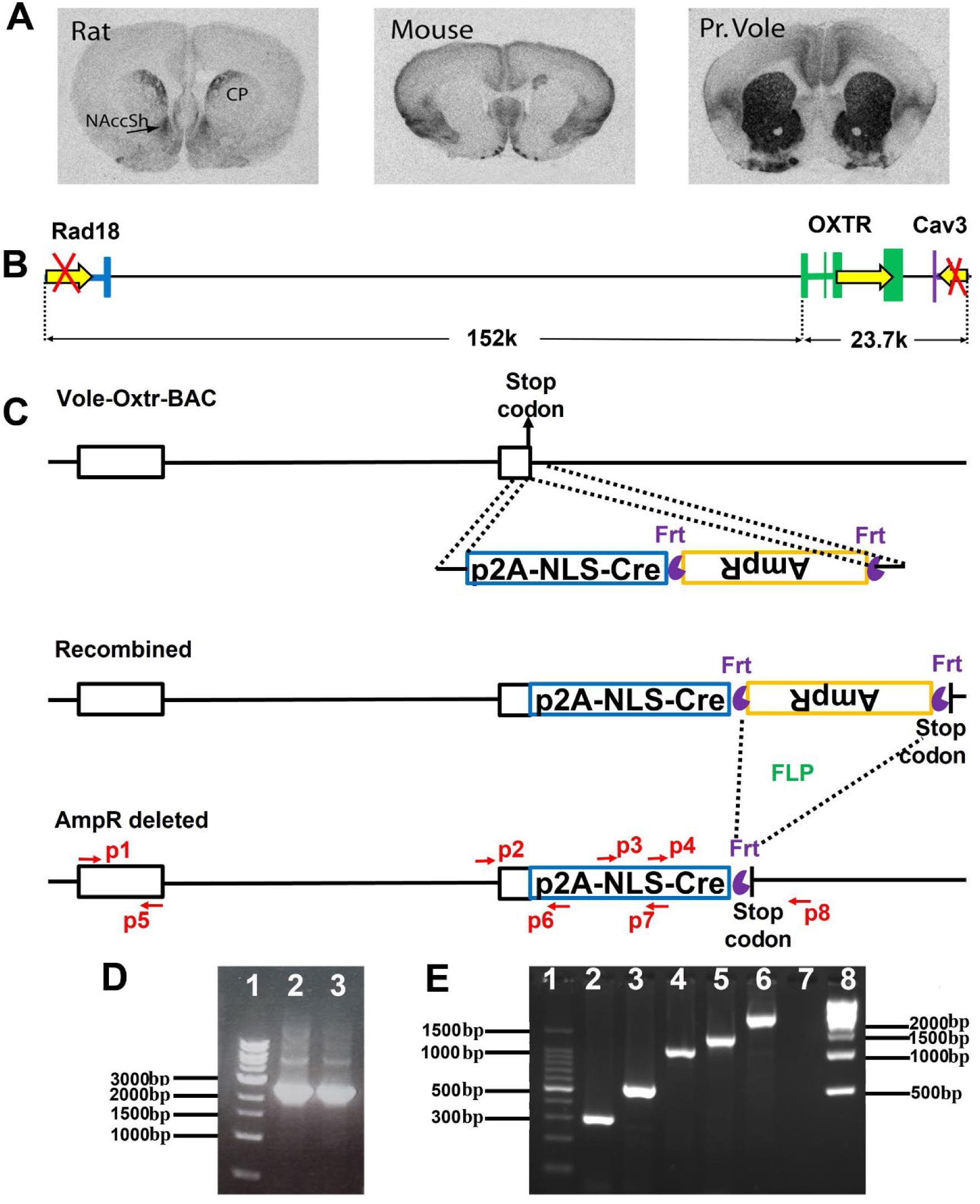
Construction of Koi lines. (A) Example of the remarkable species difference of brain OXTR distribution. OXTR receptor autoradiography in dorsal caudate putamen (CP) and nucleus accumbens shell (NAccSh) of rat, mouse, and prairie voleis (Adapted Froemke and Young(Froemke and Young, 2021)). (B) The pv*Oxtr* BAC clone contains the entire sequence of pv*Oxtr* gene (colored green), the complete intergenic sequence both upstream and downstream of <u>pv</u>*Oxtr* gene, the 12^th^ exon of *Rad18* (colored blue) and the 2^nd^ exon of *Cav3* (colored purple). The yellow arrows indicate the direction of transcription. (C) Schematic diagram of the strategy to generate Koi mice. The P2A-NLS-Cre-Frt-Amp-Frt cassette was inserted in-frame right before the stop codon of *Oxtr*. And the ampicillin selection marker was deleted by 706-Flpe. PCR primers for verifying the targeted alleles are shown as red arrows. (D) Two correctly modified vector clones were verified by using PCR with p2 and p8. (E) Koi lines were verified by 5 pairs of primers. From Lane1 to Lane8 are 100bp marker, p3+p7, p2+p6, p2+p7, p4+p8, p2+p8, blank and 1kb marker.

### Each Koi line showed a unique expression pattern of transgene in brain, yet conserved expression in mammary gland

We systemically evaluated 8 Koi lines by crossing them with ROSA-26-NLS-*LacZ* mice, a Cre reporter mouse line expressing nuclear-localized beta-galactosidase (Zhang et al., 2016; Kandasamy et al., 2021). Since Lac-Z is only expressed in the nucleus of double positive (CRE+, LacZ+) cells, but not in the neuronal processes, we could easily assess the expression pattern of our transgene. We found that each Koi founder line displayed a unique pattern of expression in brain distinct from that of wildtype mice or prairie voles (Fig.2 A-O, and Supplementary Fig.2). Almost all Koi lines showed transgene expression in olfactory bulb, lateral septum (Fig.2 F-I and Supplementary Fig.2 B,E,H,K) and ventromedial nucleus of the hypothalamus (Fig.2 K-N and Supplementary Fig.2 C,F,I,L), though with different intensities, suggesting that these regions constitute a stable core *Oxtr* expression network, as these regions also express OXTR in mice and voles (Newmaster et al., 2020; Inoue et al., 2022). Several Koi lines expressed the transgene in some brain regions specific for vole *Oxtr* expression, including nucleus accumbens (NAc) (Fig.2 A,B,D and Supplementary Fig.2 A,G,J), prefrontal cortex (PFC) (Fig.2 A,B,C and Supplementary Fig.2 J), lateral amygdala (Fig.2 K,L,M and Supplementary Fig.2 C,I,L) and deep layers of cingulate cortex (Fig.2A,B,C and Supplementary Fig.2 C,L), which suggested that cis regulatory elements in the BAC are capable of mediating the expression of *Oxtr* in the reward and reinforcement circuitry of brain, albeit depending on the sequence/structure of distal sequences >150 kb upstream and >21 kb downstream of the transcription start site. None of the Koi lines exactly “mirrored” the vole-specific expression patten. Six of the Koi lines expressed the transgene in broad areas of thalamus (Fig.2 K,L,N and Supplementary Fig.2 C,I,L), reminiscent of the strong expression of V1a vasopressin receptor (*Avpr1a*) in the thalamus (Young and Zhang, 2021). Intriguingly, the expression of OXTR in mammary glands showed similar pattern among all the Koi lines and *mouseOxtr(mOxtr)-Ires-Cre* knock-in line (Fig.2 P-T and Supplementary Fig.3).

**Fig. 2.**
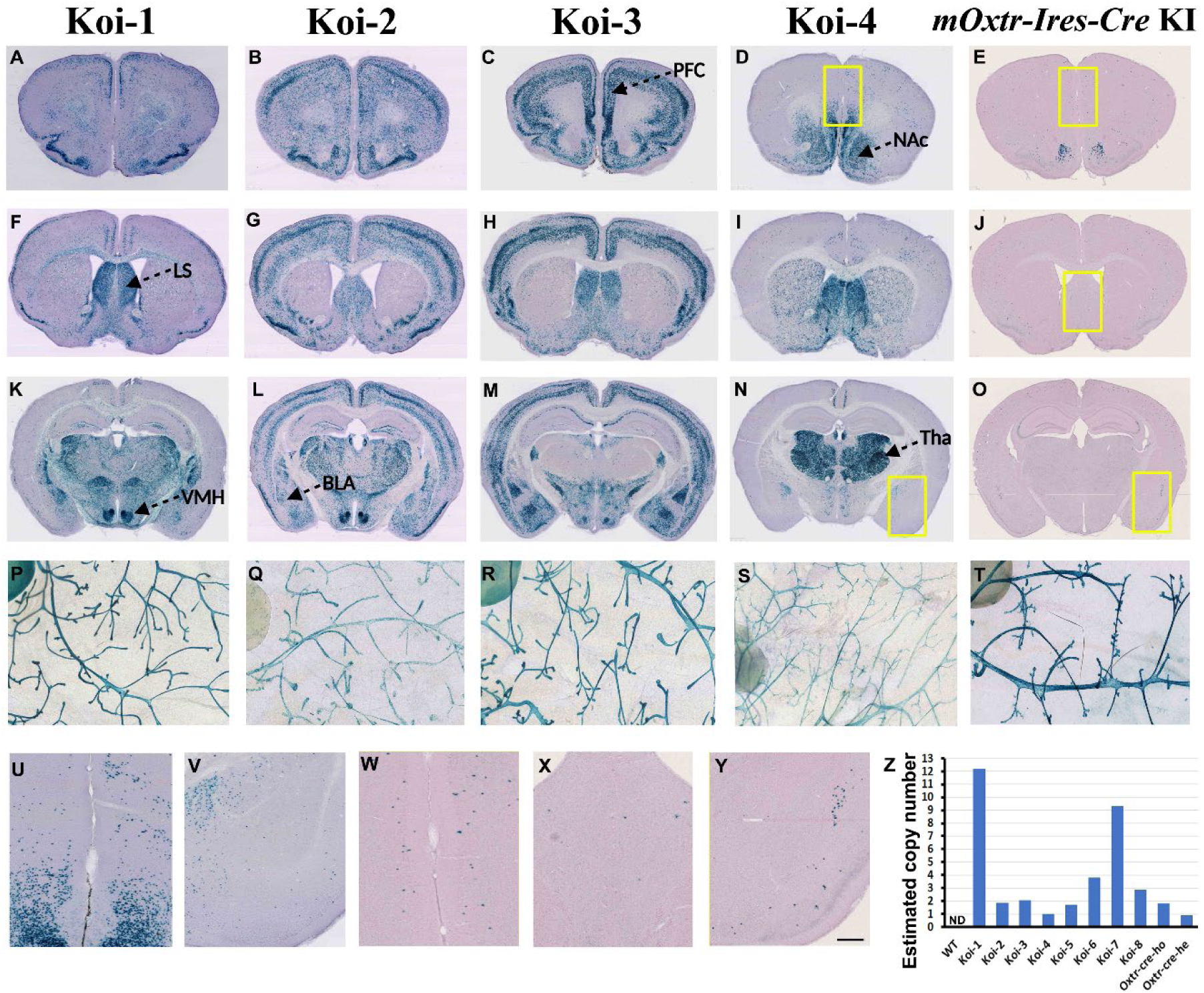
Transgenic CRE mediated Lac-Z reporter gene expression in the brain and mammary gland of Koi lines. X-gal staining of the brains and mammary glands are from the double positive (*Cre*+, *LacZ*+) offspring from Koi-1 (A, F, K, P), Koi-2 (B, G, L, Q), Koi-3 (G, H, M, R), Koi-4 (D, I, N, S) and *mOxtr-Ires-Cre* knock-in line (E, J, O, T). The *mOxtr-Ires-Cre* KI line used contains Cre (and an HA tag) inserted in the third exon, right after the start codon. Note that regardless of the insertion site of Cre in the mouse genome DNA or pvOxtr-BAC transgene, *Oxtr* expression visualized by Lac-Z staining in the mammary gland was comparable between the mOxtr-Ires-Cre KI and Koi lines. This contrasts with the diverse expression patterns of *Oxtr* expression in the brain. U, V, M, X, Y are the magnified images of the areas within the yellow-border squares from D, N, E, J, O, respectively, to show the distribution of low-density Lac-Z positive cells in these areas. Scale bar=1mm (A-O), 500um (P-T) and 200um (U-Y). (Z) The estimated copy number of transgene of all Koi lines. See Supplementary Figure 2 for images of Koi-5-8 lines. PFC=prefrontal cortex, LS=lateral septum, BLA=basolateral amygdala, VMH=ventromedial nucleus of the hypothalamus, Tha=thalamus.

Transgenes that are introduced by pronuclear injection typically integrate into a single site of the genome as tandem concatemers (Smirnov et al., 2019). BAC copy number variation and different insertional loci can lead to distinct patterns of ectopic expression, and increased BAC transgene copy numbers often correlate with increased BAC gene expression (Gong et al., 2007). We then examined the relationship between transgene copy number and LacZ signal using quantitative genomic PCR (Chandler et al., 2007). As expected, the copy number of WT, heterozygous and homozygous *mOxtr-Ires-Cre* knock-in mice was 0, 1 and 2 respectively (Figure.2 Z). The copy number of transgene in Koi lines are: Koi-1 (12), Koi-2 (2), Koi-3 (2), Koi-4 (1), Koi-5 (2), Koi-6 (4), Koi-7 (9), Koi-8 (3). Koi-6 and Koi-7 showed the most limited expression of reporter gene, though the copy number of transgene were 4 and 9 respectively. Koi-4 showed strong expression of reporter gene in NAc, septum and thalamus, though there was only 1 copy of transgene. Therefore, most likely integration site rather than copy number is the reason of the variant BAC transgene expression patterns across our Koi lines.

### Adult brain OXTR binding patterns differ across Koi lines

To detect the distribution of OXTR protein in adult Koi lines without the interference of endogenous mouse OXTR (mOXTR) signal, Koi lines were crossed with *mOxtr^−/−^* line twice sequentially to produce Koi: m*Oxtr*^−/−^ mice, and pvOXTR binding was detected using receptor autoradiography. We focused on analyzing 4 lines: Koi-1: m*Oxtr*^−/−^, Koi-2: m*Oxtr*^−/−^, Koi-3: m*Oxtr*^−/−^ and Koi-4: m*Oxtr*^−/−^ based on the strong expression of reporter gene in behaviorally relevant neuronal populations. Autoradiography demonstrated functional pvOXTR in brain resembling to some extent Cre-induced reporter gene expression (Fig.3) and OXTR expression in prairie vole (Supplementary Fig.4). Brain OXTR binding of Koi-3: m*Oxtr*^−/−^ was consistent with Lac-Z expression in the reporter line (Fig.2 C, H, M and Figure.3 C, H, M). Interestingly, the OXTR binding in PFC of Koi-3: m*Oxtr*^−/−^ was similar to that of prairie vole (Figure.3 C). In contrast, Koi-4: m*Oxtr*^−/−^ showed low OXTR binding in NAc when compared with the high Lac-Z staining (Fig.2 D and Figure.3 D). Koi-2: m*Oxtr*^−/−^, the line which displayed the expression of reporter genes in the broadest regions, did not show strong signal in autoradiography. It is interesting that Layer IV of cortex in Koi-4: m*Oxtr*^−/−^ showed strong OXTR binding (Figure.3 I and N), which is highly likely to originate from the axon projections from the thalamus (Fig.2 N), as was previously reported for NAc projections in voles (Inoue et al., 2022). Consistent with Lac-Z expression, we do not see a clear relationship between OXTR binding and transgene copy number. For example, Koi-1: m*Oxtr*^−/−^ showed rather low OXTR signal (Fig.3 A, F, K) with the highest copy number, while Koi-4: m*Oxtr*^−/−^ showed very strong OXTR signal at lateral septum and thalamus (Fig.3 I, N), with the lowest copy number.

**Fig. 3.**
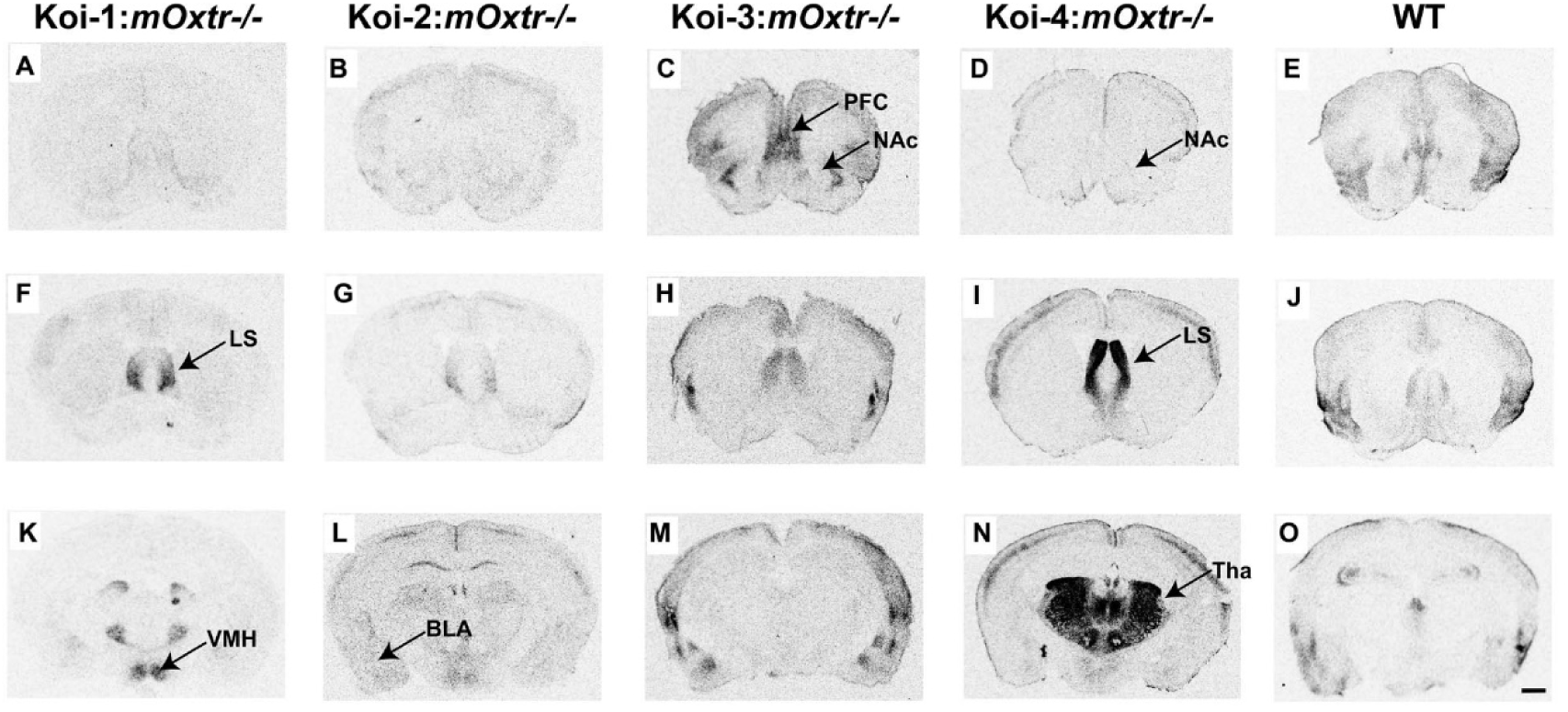
The expression of OXTR in adult brain of Koi lines. Autoradiographs illustrate the distribution of OXTR binding in adult Koi-1: m*Oxtr*^−/−^ (A, F, K), Koi-2: m*Oxtr*^−/−^ (B, G, L), Koi-3: m*Oxtr*^−/−^ (C, H, M), Koi-4: m*Oxtr*^−/−^ (D, I, N) and wild type (WT) (E, J, O). All Koi mice used for this analysis were pv*Oxtr-P2A-Cre*: m*Oxtr*^−/−^ mice in which the pvOXTR was expressed while endogenous mOXTR was absent. PFC=prefrontal cortex, LS=lateral septum, BLA=basolateral amgydala, VMH=ventromedial nucleus of the hypothalamus, Tha=thalamus. Scale bar=1mm.

*Oxtr* mRNA level in Koi-3: m*Oxtr*^−/−^ and Koi-4: m*Oxtr*^−/−^ was further investigated with Fluorescent RNAscope *in situ* hybridization (Fig.4). Consistent with Lac-Z expression in the reporter line, Koi-3: m*Oxtr*^−/−^ showed strong signal in PFC and Ctx (Fig.4 A, C), while Koi-4: m*Oxtr*^−/−^ showed prominent signal in NAc and Thalamus (Fig.4 B, D). qRT-PCR analysis further revealed that adult Koi-4: m*Oxtr*^−/−^ mice had the highest Oxtr mRNA signal in the ventral striatum, several fold higher than WT, consistent with the Lac-Z staining in the reporter line (Fig.5 A, B). The discrepancy of OXTR signal in NAc of Koi-4: m*Oxtr*^−/−^ between autoradiography and Lac-Z staining /RNAscope/qRT-PCR may, therefore, be due to the translational regulation or differences in the sensitivity of the detecting methods. In addition, qRT-PCR analysis revealed no significant differences in *Oxtr* expression levels in the mammary gland among the different Koi lines (Supplementary Fig.5).

**Fig. 4.**
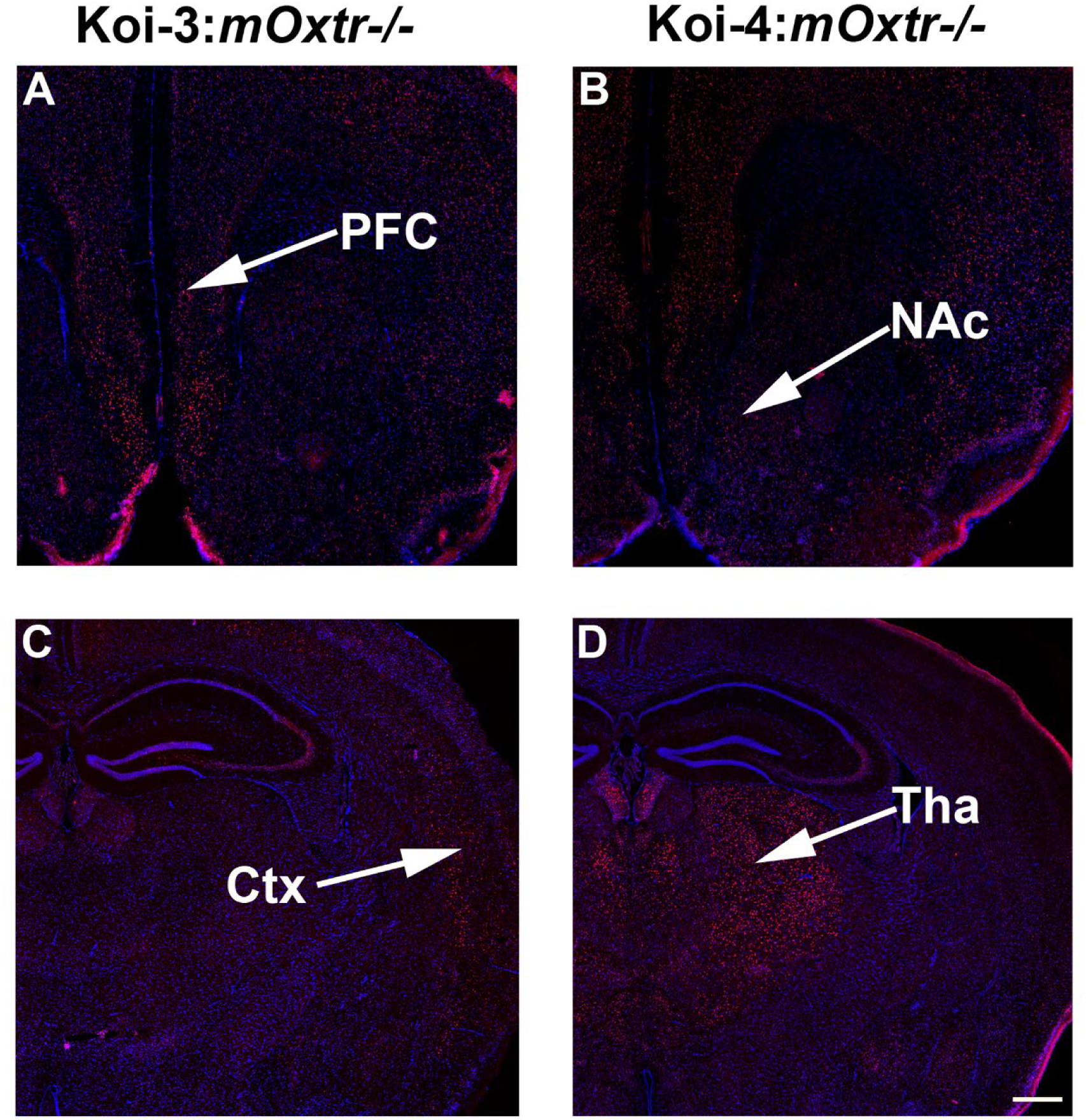
mRNA expression of *Oxtr* in the adult brain of Koi lines. *Oxtr* mRNA (red) was labeled with in situ hybridization against DAPI for nuclei labelling (purple). (A) and (C) show clear expression of *Oxtr* in the PFA and Ctx in Koi-3: m*Oxtr*^−/−^. (B) and(D) show clear expression of *Oxtr* in the NAc and Tha in Koi-4: m*Oxtr*^−/−^. All Koi mice used for this analysis were pv*Oxtr-P2A-Cre*: m*Oxtr*^−/−^ mice, in which the pvOXTR was expressed while endogenous mOXTR was absent. PFC=prefrontal cortex, NAc= Nucleus accumbens, Ctx=Cortex, Tha=thalamus. Scale bar=400um for A and B, 500um for C and D.

**Fig. 5.**
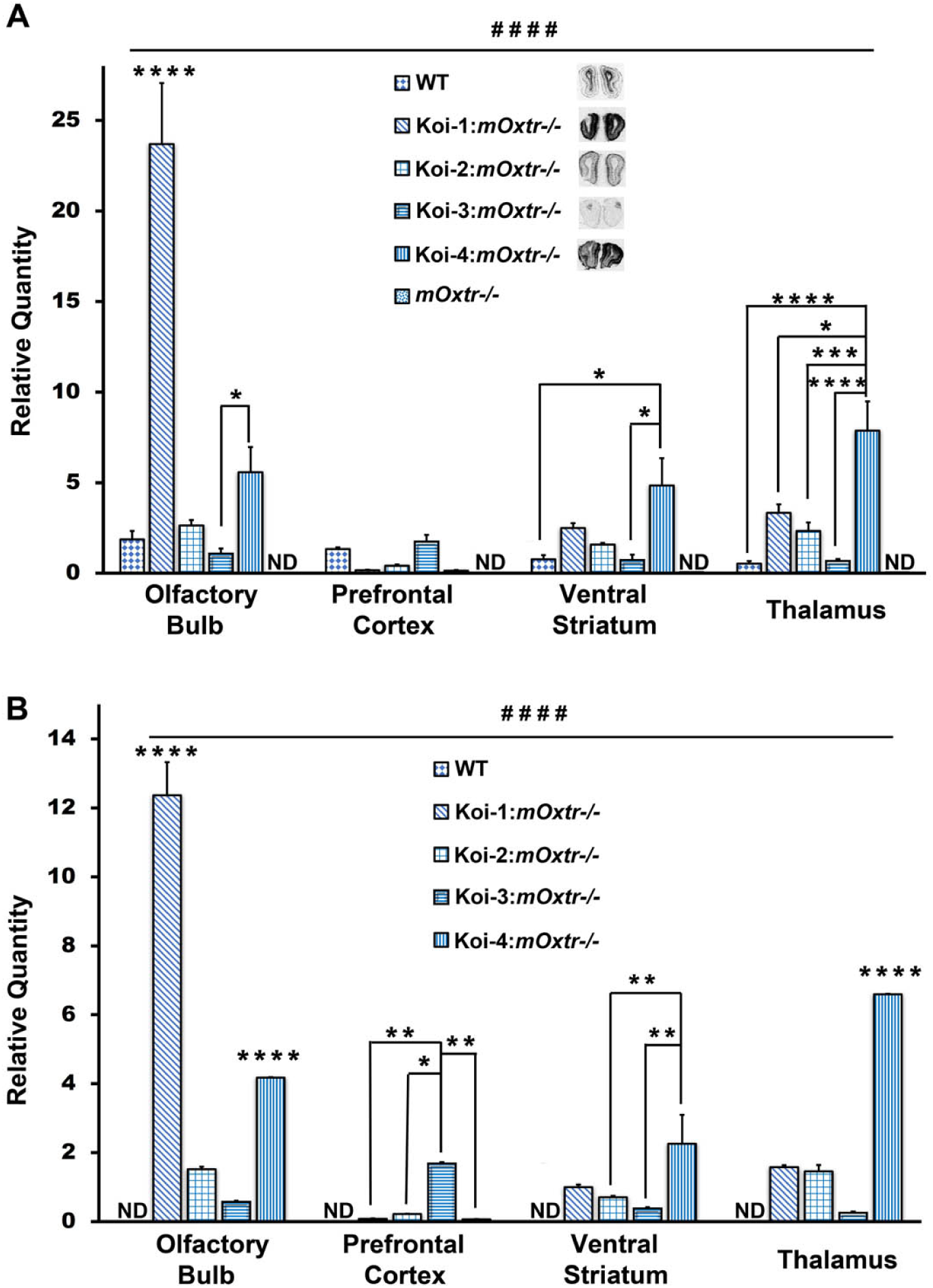
Quantitative PCR analysis of *Oxtr* mRNA expression in different brain regions of Koi lines. **(A)** A primer set recognizing both m*Oxtr* and pv*Oxtr* was used to analyze *Oxtr* mRNA in olfactory bulb, prefrontal cortex, ventral striatum and thalamus from WT, Koi-1: m*Oxtr*^−/−^, Koi-2: m*Oxtr*^−/−^, Koi-3: m*Oxtr*^−/−^, Koi-4: m*Oxtr*^−/−^ and *mOxtr^−/−^* mutant mice, n=4 for each genotype. Note that the level of *Oxtr* mRNA in *mOxtr^−/−^* mutant mice was nondetectable (ND). The autoradiograph of OXTR binding in olfactory bulb from each line were shown alongside the legend. There was a significant interaction between position and genotype F (12,64) =26.33, p<0.0001; Post hoc Bonferroni test revealed that *Oxtr* expression level in olfactory bulb of Koi-1: m*Oxtr*^−/−^ was significantly higher than that of other genotype groups (p<0.0001 for all), and that of Koi-4: m*Oxtr*^−/−^ was significantly higher than that of Koi-3: m*Oxtr*^−/−^ p<0.05. And the striatum of Koi-4: m*Oxtr*^−/−^ expressed significantly higher *Oxtr* when compared with WT and Koi-3: m*Oxtr*^−/−^ (p<0.05 for both). The *Oxtr* expression level in thalamus of Koi-4: m*Oxtr*^−/−^ was significantly higher than other genotype groups (p<0.0001 when compared with WT and Koi-3: m*Oxtr*^−/−^, p<0.001 when compared with Koi-2: m*Oxtr*^−/−^, p<0.05 when compared with Koi-1: m*Oxtr*^−/−^) (B) A primer set specifically recognizing pv*Oxtr.* was used to analyze *Oxtr* mRNA in WT, Koi-1: m*Oxtr*^−/−^, Koi-2: m*Oxtr*^−/−^, Koi-3: m*Oxtr*^−/−^ and Koi-4: m*Oxtr*^−/−^ mic, n=4 for each genotype. Note that the level of *Oxtr* mRNA in WT mice was barely detectable. There was a significant interaction between position and genotype F (9,48) =89.25, p<0.0001; Post Hoc Bonferroni test revealed that *Oxtr* expression level in the olfactory bulb of Koi-1: m*Oxtr*^−/−^ was significantly higher than that of other genotype groups (p<0.0001 for all), and that of Koi-4: m*Oxtr*^−/−^ was significantly higher than that of Koi-2: m*Oxtr*^−/−^ and Koi-3: m*Oxtr*^−/−^ (p<0.0001 for both). The *Oxtr* expression level in prefrontal cortex of Koi-3: m*Oxtr*^−/−^ was significantly higher than that of other groups (p<0.01 when comparing with Koi-1: m*Oxtr*^−/−^ and Koi-4: m*Oxtr*^−/−^; p<0.05 when comparing with Koi-2: m*Oxtr*^−/−^). And the striatum of Koi-4: m*Oxtr*^−/−^ expressed significantly higher *Oxtr* when comparing with Koi-2: m*Oxtr*^−/−^ and Koi-3: m*Oxtr*^−/−^ (p<0.01 for both). The *Oxtr* expression level in thalamus of Koi-4: m*Oxtr*^−/−^ was significantly higher than that of other genotype groups (p<0.0001 for all). All Koi mice used for this analysis were pv*Oxtr-P2A-Cre*: m*Oxtr*^−/−^ mice, in which the pvOXTR was expressed while endogenous mOXTR was absent. The significance of interaction between genotype and brain region: #### p<0.0001. The significance of single main effect of genotype: *p<0.05; **p<0.01; ***p<0.001; ****p<0.0001.

Although none of our Koi lines mirrored the endogenous OXTR expression in prairie voles, we confirmed strong expression of pv*Oxtr* in olfactory bulb of Koi-1line, PFC and amygdala of Koi-3line, striatum and thalamus of Koi-4 lines, and the broad expression of pv*Oxtr* in Koi-2 line. As OXTR signaling in some of these regions are similar to that in voles, albeit in separate lines, and postulated to be involved in vole social behavior (Walum and Young, 2018a), we used these lines to address the question of whether variation in *Oxtr* transcription in brain can lead to variation in social behaviors.

### Different Koi lines showed diverse behaviors in a Partner Preference Test

The partner preference test is widely used as a laboratory proxy for pair bonding in prairie voles. We initially evaluated partner preference in both male and female Koi mice on a m*Oxt*^+/+^ background. However, we did not observe consistent differences in partner preference among the Koi lines in males. In contrast, we found that Koi-4 female mice showed a preference for their mate over a novel male. We therefore focused our subsequent analyses on females. Again, we bred these Koi lines with *mOxtr^−/−^* mice (both on C57/B6 background) to generate Koi: m*Oxtr*^−/−^ offspring expressing only pv*Oxtr*. PPTs were performed in ovariectomized females after 21 days of cohabitation with 3 mating bouts elicited by estrogen and progesterone injections on days 2, 9 and 16. The PPT consisted of three phases (Fig. 6A, B). During the habituation phase, the experimental female was habituated to the chamber for 5 minutes with both pen-boxes empty. During the recognition phase, the partner male was restricted in one pen-box, and a novel “stranger” male was restricted in another pen-box, the experimental female was allowed to explore the arena for 10 minutes to test social novelty preference. During the preference phase, the positions of the partner and stranger were switched to avoid position bias, and the female was allowed to explore the chamber for an additional 20 minutes. The stimulus animals were wild-type mice to minimize potential confounding effects arising from genotype-dependent interactions between stimulus and focal animals.

**Fig. 6.**
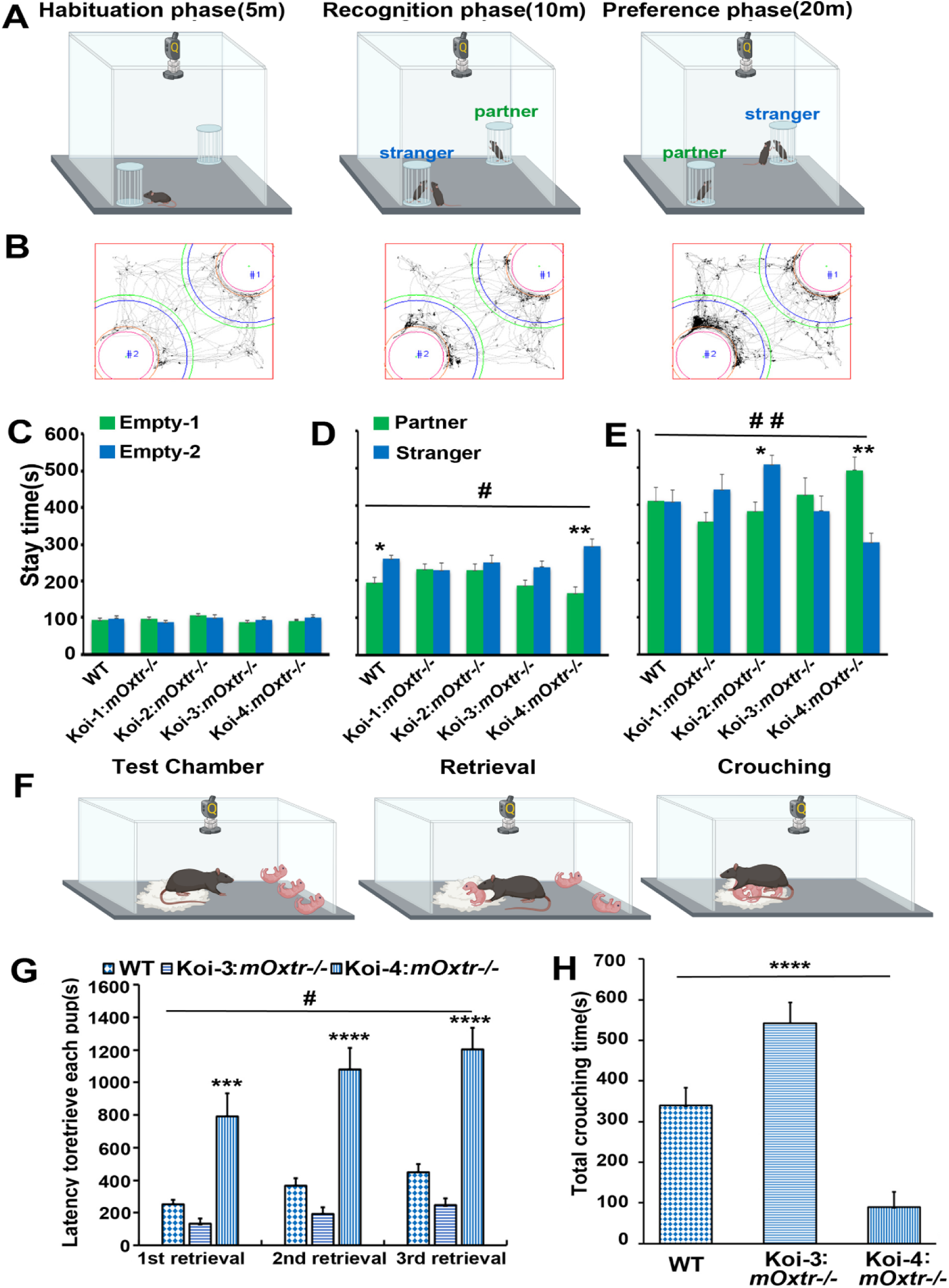
Social behaviors of Koi lines. (A) Illustration of the partner preference test (PPT) paradigm. The experimental mouse was placed in a chamber with an automatic monitoring system, where it could freely explore two pen-boxes localized in two diagonal corners. Habituation phase: two pen-boxes are empty. Recognition phase: a stimulating partner mouse is placed into one pen-box and a stimulating stranger mouse is placed in the other one. Preference phase: The positions of two stimulating mice are switched. (B) Representative traces of one experimental mouse during the three phases. (C-E) The stay time in the surrounding area of each pen box during three phases;(C) Habituation phase;(D) Recognition phase; (E) Preference phase. Samples sizes for the PPT are wild type(WT) (n=12), Koi-1: m*Oxtr*^−/−^ (n=14), Koi-2: m*Oxtr*^−/−^ (n=13) Koi-3: m*Oxtr*^−/−^ (n=12) Koi-4 : m*Oxtr*^−/−^ (n=14). All the experimental mice are female. (F) The illustration of the procedure of the maternal behavior test. (G) Pup retrieval latency and (H) crouching time were recorded and analyzed. Sample sizes for the maternal behavior tests are WT (n=15), Koi-3: m*Oxtr*^−/−^ (n=18), Koi-4: m*Oxtr*^−/−^ (n=18). Significance of interaction between independent factors: # p<0.05; ## p<0.01. Significance of main effect or single main effect of genotype: *p<0.05; **p<0.01; ***p<0.001; ****p<0.0001.(Illustrations created with Biorender.com). All Koi mice used for this analysis were pv*Oxtr-P2A-Cre*: m*Oxtr*^−/−^ mice in which the pvOXTR was expressed while endogenous mOXTR was absent.

Mixed ANOVA was conducted to examine the effect of genotype and stimulus animal (pen-box position during habituation) on the area stay time. In the habituation phase (Fig. 6C), there was no significant genotype × position interaction (F(4, 60) = 0.69, p = 0.60), and there was no significant main effect of position (F(1, 60) = 0.03, p = 0.87), indicating no position preference for all Koi lines.

In the recognition phase (Fig.6 D), WT and Koi-4: *mOxtr^−/−^*mice spent significantly more time in the stranger area than in the partner area, whereas Koi-1, Koi-2, and Koi-3: *mOxtr^−/−^* mice did not show a significant preference for either stimulus animal. There was a significant main effect of stimulus animal (F (1,60) =15.38, p=0.0003), a main effect of genotype F (4,60) =2.62, p=0.04, and a significant genotype × stimulus animal interaction F (4, 60) =2.61, p=0.04. Bonferroni-adjusted pairwise comparisons showed that Koi-4: *mOxtr^−/−^* and WT mice spent significantly more time in the stranger area than in the partner area (p = 0.0001 and p = 0.028, respectively), demonstrating a preference for social novelty. No significant difference between the time spent in the “stranger” and “partner” areas was observed for Koi-1: m*Oxtr*^−/−^, Koi-2: m*Oxtr*^−/−^, or Koi-3: m*Oxtr*^−/−^ mice (p = 0.96, 0.49, and 0.09, respectively).

In the preference phase (Fig. 6E), after the two stimulus animals had presumably become familiar during the initial 10-min recognition phase, Koi-4: *mOxtr^−/−^* mice spent significantly more time in the partner area than in the stranger area, whereas Koi-2: *mOxtr^−/−^* mice spent significantly more time in the stranger area than in the partner area. WT, Koi-1: *mOxtr^−/−^*, and Koi-3: *mOxtr^−/−^* mice showed no significant preference for either stimulus animal. There was no significant main effect of stimulus animal (F(1, 60) = 0.04, p = 0.84) or genotype (F(4, 60) = 1.36, p = 0.26), but there was a significant genotype × stimulus animal interaction (F(4, 60) = 4.64, p = 0.003). Bonferroni-adjusted pairwise comparisons showed that Koi-2: *mOxtr^−/−^*mice spent significantly more time in the stranger area than in the partner area (p = 0.024), whereas Koi-4: *mOxtr^−/−^* mice spent significantly more time in the partner area than in the stranger area (p = 0.002). No significant difference was observed for WT, Koi-1: *mOxtr^−/−^*, or Koi-3: *mOxtr^−/−^* mice (p = 0.99, 0.16, and 0.45, respectively).

### Different Koi lines showed diverse behaviors in Pup Retrieval Test

Based on the abundant pvOXTR expression in reward-related brain regions in Koi-3 and Koi-4 mice (Fig. 3C,D,H,I,M,N), and the established role of OXTR in maternal care in mice and voles (Olazábal and Young, 2006; Marlin et al., 2015; Carcea et al., 2021), we further examined parental behaviors in virgin females by the pup retrieval test from these two lines (Fig. 6F–H). The wild-type pups were chosen as stimulus animals to minimize potential confounding effects arising from genotype-dependent interactions between stimulus and focal animals.

In the pup retrieval test (Fig. 6F, G), Koi-4: m*Oxtr*^−/−^mice showed prolonged retrieval latencies compared with WT and Koi-3: m*Oxtr*^−/−^mice, whereas Koi-3: m*Oxtr*^−/−^mice showed shorter retrieval latencies than WT mice. A mixed ANOVA revealed a significant main effect of genotype on retrieval latency (F(2, 48) = 29.58, p < 0.0001). There was also a significant genotype × pup number interaction (F(2.40, 57.49) = 4.45, p = 0.011).

Bonferroni-adjusted pairwise comparisons showed that Koi-4: m*Oxtr*^−/−^mice required significantly longer to retrieve the first, second, and third pups than WT mice (p = 0.0003, p < 0.0001, and p < 0.0001, respectively) and Koi-3: m*Oxtr*^−/−^mice (p = 0.0005, p < 0.0001, and p < 0.0001, respectively). Koi-3: m*Oxtr*^−/−^mice, in turn, required significantly less time to retrieve the first, second, and third pups than WT mice (p = 0.010, 0.006, and 0.003, respectively).

Crouching behavior was also quantified (Fig. 6F, H). There was a significant effect of genotype on crouching time (one-way ANOVA, F(2, 48) = 27.56, p < 0.0001). Bonferroni-adjusted pairwise comparisons showed that Koi-3: m*Oxtr*^−/−^mice spent significantly more time crouching than WT mice (p = 0.008) and Koi-4: m*Oxtr*^−/−^mice (p < 0.0001), whereas Koi-4: m*Oxtr*^−/−^mice spent significantly less time crouching than WT mice (p = 0.001). Thus, Koi-3 and Koi-4 mice, which differ in their pvOXTR expression patterns, exhibited distinct quantitative profiles of parental behaviors.

### The TAD structure of brain *Oxtr* is markedly distinct from that of both *Oxt* and peripheral *Oxtr*

The divergent brain expression patterns across our Koi lines suggest that distal elements external to our BAC cassette contribute to the variation in *Oxtr* expression. TADs represent a key feature of hierarchical genome organization comprising regions of high chromatin inter-connectivity, and TAD formation appears to be linked to transcription regulation (Dekker et al., 2002; Dixon et al., 2012; Beagan and Phillips-Cremins, 2020; Rajderkar et al., 2023). TADs are generally conserved within the same tissue type across different species. We therefore utilized publicly available 3D chromatin datasets from mouse (Extended data Table7-1) and analyzed the TAD structure of *Oxtr* and *Oxt*, both in brain related tissues/cells and peripheral tissues/cells. We discovered that *Oxtr* is consistently located at the boundary region of TAD structures across several tissues and cell lines (Fig.7 A). Specifically, only in brain related samples (whole brain tissue, neural progenitor cells, and cortex tissue), *Oxtr* is situated at the boundary of a large inter-TAD interaction between two neighboring TADs and there existed a significant long-range interaction (Fig.7 A). In contrast, the oxytocin peptide gene, *Oxt*, which is expressed in a highly conserved brain pattern across vertebrates, lacks organized TAD structures (Fig.7 B). Previously no Hi-C dataset from mammary gland tissue of any species was available in public repositories. After we submitted our manuscript, although no mouse mammary-gland Hi-C dataset has yet become publicly available, a human breast-tissue Hi-C dataset was published (Choppavarapu et al., 2025). We therefore analyzed the Hi-C data generated from normal human mammary gland samples. Consistent with our observations in the mouse peripheral tissue/cells, the *OXTR* locus is positioned within a pronounced TAD-boundary and exhibits no detectable long-range chromatin interactions (Supplementary Fig. 6A). In contrast, the *OXT* locus lacks a canonical TAD architecture, with no obvious TAD organization observed (Supplementary Fig. 6B). These findings demonstrate that the TAD landscape differ dramatically between *Oxtr* and *Oxt*, and highlight that the 3D genomic architecture surrounding the *Oxtr* gene varies between brain tissues/cells and peripheral tissue/cells. There exists clear long-distance interaction in brain *Oxtr* gene TAD structure. The presence of extensive long-distance chromatin interactions around *Oxtr* in brain tissues may at least partially explain why its expression is hypersensitive to distal regulatory sequences beyond the range of a BAC construct. Moreover, such long-distance interactions likely expose *Oxtr* regulation to a broader set of cis- and trans-acting influences, rendering brain *Oxtr* expression inherently more variable across individuals and species.

**Fig. 7.**
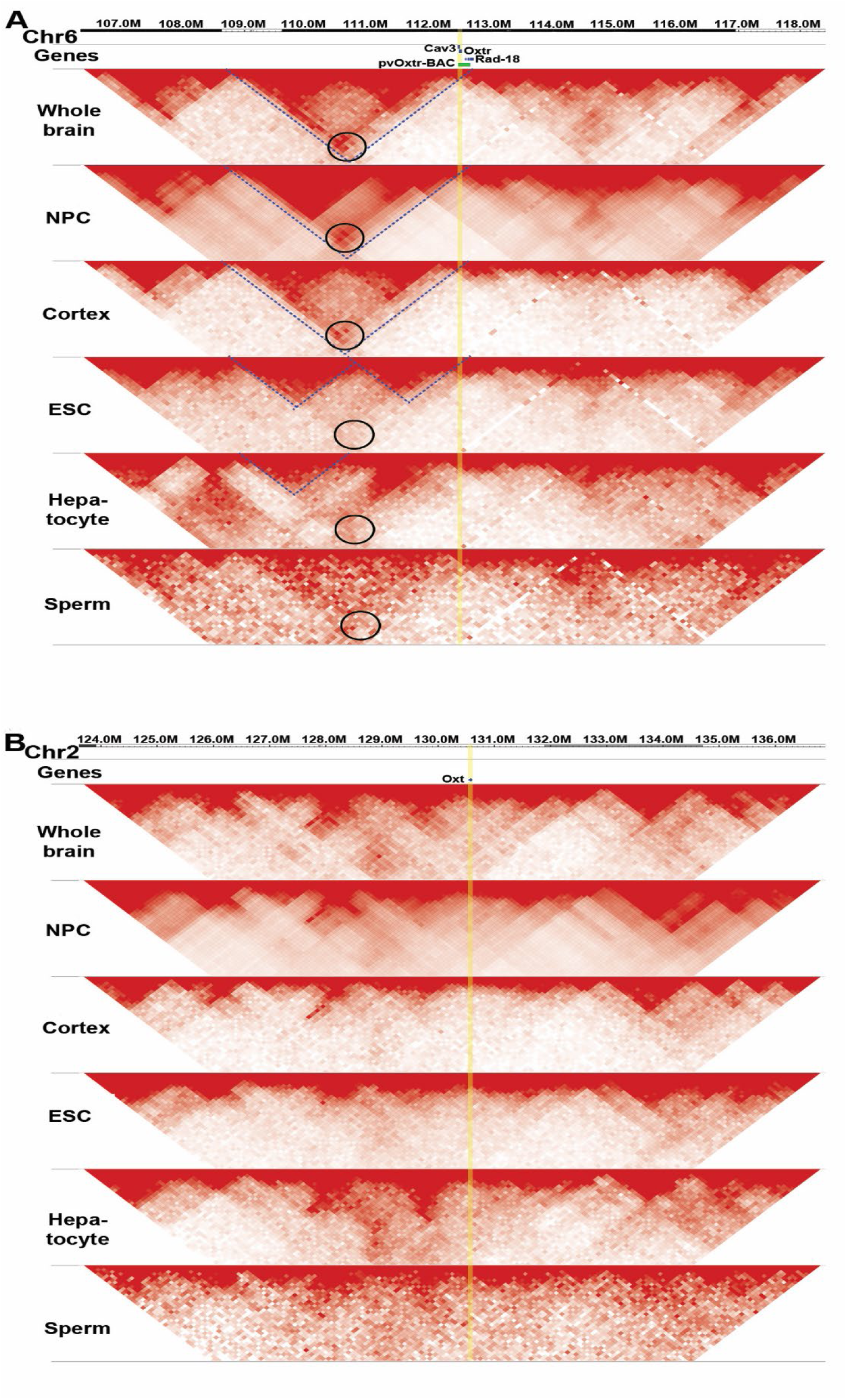
Heatmap of the chromatin contacts surrounding *Oxtr* and *Oxt* across mouse tissues/cells from megabase-size visualization. (A) Heatmap of the chromatin contacts surrounding *Oxtr* across mouse tissues/cells: *Oxtr* is located at a TAD boundary region. The increase of interactions characterizes the TADs structure and the interaction of two neighboring TADs can be more clearly observed in brain-related samples. Blue dashed lines were prepared to help observing the TAD structures, and black circles were prepared to help observing the long-distance interaction obtained from Hi-C interaction matrices surrounding *Oxtr*. (B) Heatmap of the chromatin contacts surrounding *Oxt* across mouse tissues/cells. No TAD structure was observed surrounding *Oxt* locus.

## Discussion

### Diversity of *Oxtr* expression patterns causes diversity of social behaviors

The diverse OXTR expression of our Koi lines is reminiscent of the diverse OXTR expression across different species. The unique expression pattern of *Oxtr* in monogamous prairie vole was mainly observed in PFC, NAc and BLA. We obtained expression in all of these specific regions, but in 3 separate lines. These Koi lines with distinct brain OXTR expression pattern provided an opportunity to test the effect of variation of OXTR expression on diverse social behaviors.

It is intriguing that Koi-4 line showed preference to their mating partners but not strangers, which is a behavior specific in prairie voles but not in mice. Multiple studies suggested that NAc and PFC play important roles in mediating partner preference behavior in prairie voles (Amadei et al., 2017; Walum and Young, 2018b). Consistently, the Koi-4 line showed robust transgene expression in these brain regions. Notably Koi-4 line also demonstrated strong *Oxtr* expression in the entire thalamus region, which reminds us the distribution of *Avpr1a* (Phelps and Young, 2003). Given the fact that OXTR and AVPR1A genes were derived from a duplication of a single ancestral gene (Grinevich et al., 2016), it is not surprising that the similar expression regulation machinery exists. Recent studies indeed revealed the critical functions of different thalamic nuclei in social memory and cognition (Rangel et al., 2018; Wolff and Vann, 2019; Kappel et al., 2022; Chen et al., 2024). Partner preference formation and maintenance need to recruit multi- and cross-modality sensation, social recognition and social memory, selective attention, arousal, and reward systems. We propose a model that the sufficient upregulation of the OXTR expression in one or multiple of these systems can tune the network and boost partner preference behavior. In addition, the maternal behavior in Koi-3 and Koi-4 lines was enhanced and reduced, respectively. This result suggested that OXTR signaling facilitates social partner preference and maternal care by coordinating activity across different neural networks. A main conclusion of our study is that simply by redistributing OXTR binding in space, variation in social preference and parental behaviors emerged, which conceivably could impart advantageous behaviors that natural selection could act upon and amplify. Future study will determine how the variation in OXTR distribution in our Koi lines affects neural circuits mediating social behaviors.

Recent work has shown that constitutive *Oxtr* deletion in prairie voles does not abolish pair bonding or parental behaviors(Berendzen et al., 2023). This finding may reflect an important distinction between the complete absence of OXTR signaling from development onward and variation in the spatial distribution of OXTR expression. In constitutive *Oxtr* knockout voles, the absence of OXTR signaling throughout development may allow compensatory or redundant mechanisms to emerge that preserve social behaviors. Consistent with this possibility, Young has proposed that the timing of OXTR loss may be critical, as neural circuits may become functionally dependent on oxytocin signaling through developmental exposure to this hormone. Our Koi system provides a complementary approach by preserving pvOXTR expression while generating distinct spatial expression patterns within the same mouse genetic background. The divergent social phenotypes observed across the Koi lines suggest that variation in the spatial organization of OXTR expression, rather than simply the presence or absence of OXTR signaling, may contribute to behavioral diversity. Thus, the preservation of social behaviors in constitutive *Oxtr* knockout voles does not preclude a role for OXTR expression patterns in shaping behavioral variation.

### A potential genetic mechanism contributing to diversity of brain OXTR expression

The sex steroid and oxytocin systems play important roles in modulating reproductive and related social behaviors in vertebrates. Sexual behavior, e.g., the motivation to mate and motor patterns, is highly conserved across vertebrate species. By contrast, social behaviors, e.g., sociality, mating strategy (monogamy vs polygamy), parental and alloparental behaviors are more nuanced and can vary dramatically across species in the same genus or even between individuals of a single species. It is interesting, therefore, that the distribution of steroid receptors is quite conserved among species (Young and Crews, 1995), while that of the OXTR is extremely diverse across and even within species (Froemke and Young, 2021; Young and Zhang, 2021). It is even more interesting that although the distribution of OXTR is diverse, its ligand, oxytocin, is expressed in a highly conserved neuroanatomical pattern in vertebrates (Ross and Young, 2009). Previous work showed that three independent rat lines with puffer fish oxytocin BAC transgene (containing 16 kb of 5’ flanking regulatory sequence) expressed the *Fugu* oxytocin transcript specifically in oxytocin neurons in rat (Venkatesh et al., 1997). Moreover, a similar transgenic experiment transferring approximately 5 kb of the *Fugu* oxytocin already resulted in a faithful expression of the fish genes in mouse oxytocin neurons (Gilligan et al., 2003). These studies suggest there is a remarkably conserved transcriptional regulatory machinery across species, even between the *Fugu* and rodent oxytocin genes, despite being separated by 400 million years of evolution.

The GENSAT (Gene Expression Nervous System Atlas) Project demonstrated that most BAC transgenes reliably recapitulate endogenous expression across independent lines, provided that extensive flanking intergenic regions are included (Gong et al., 2003; Heintz, 2004; Gong et al., 2007; Gerfen et al., 2013; Schmidt et al., 2013). Our pv*Oxtr* clone contained the entire locus with full upstream and downstream intergenic sequences, yet the transgenic lines we generated displayed strikingly divergent brain expression patterns. These differences were not explained by transgene copy number, but instead reflected strong insertion-site effects. This paradox contrasts with the general reproducibility of BAC transgenics and echoes the observation that virtually every rodent or primate species has a unique distribution of OXTR binding in the brain.

The 3D genomic structure analysis showed that the *Oxtr* loci sit within large TAD structures, increasing its exposure to distal regulatory elements through an expanded interaction landscape, whereas *Oxt* lacks organized TAD structures. This difference might explain why short promoter sequences suffice for conserved *Oxt* expression across species, while even a large BAC is insufficient for precise, reproducible transcription regulation of *Oxtr*. The large TAD and extensive long-range interactions render *Oxtr* especially sensitive to chromosomal context, accounting for the position effects we observed.

It is also noteworthy that pv*Oxtr* expression in mammary gland (which is responsible for essential lactation in mammals) was conserved across all Koi lines, indicating that the essential cis-regulatory elements for peripheral expression are captured within the BAC and are resistant to position effects. This conclusion is consistent with the chromatin architecture we observed: the chromatin landscape differs considerably between brain-related tissues and peripheral tissues. There exists an obvious increase of long-distance interactions between neighboring TADs, forming a larger TAD structure in brain-related tissues but not peripheral tissues. Moreover, critical peripheral regulatory sequences may be clustered proximally around the locus, so that they fall entirely within the BAC. By comparison, in brain tissues, regulatory elements are dispersed across a much broader genomic interval, extending beyond the BAC. Together, the larger interaction landscape and distributed regulatory architecture in brain tissues likely explain why BAC constructs are sufficient for peripheral but not brain expression of *Oxtr*. Our TAD analyses provide a correlative framework supporting the involvement of distal regulatory architecture in shaping brain-specific *Oxtr* expression, but do not establish a causal relationship. Demonstrating a causal role for TAD organization in distal *Oxtr* regulation will require direct perturbation of chromatin architecture, such as targeted disruption of TAD boundaries or chromatin looping interactions using genome-engineering approaches in the future.

Our discovery is well aligned with broader principles of genome organization: subtle changes within 3D chromatin structure may result in substantial changes in local gene expression (Xiao et al., 2021). TADs also display increased SNPs and structure variations (SVs) density and higher recombination rate compared to inter-TAD regions(Dekker et al., 2002; Dixon et al., 2012; Beagan and Phillips-Cremins, 2020; Rajderkar et al., 2023). In fact, SNPs in the *Oxtr* intron explains 74% of the variance in striatal *Oxtr* expression and social attachment in prairie voles (King et al., 2016) and it is possible that these SNPs near *Oxtr* are linked to variations of long-range chromosomal interaction. Collectively, these evidences support the notion that brain *Oxtr* expression is hyper-sensitive to genetic variation.

Such an organization may endow *Oxtr* with enhanced evolvability. Oxytocin is a secreted neuromodulator that can diffuse along axons and through the extracellular space(Leng and Ludwig, 2008; Mitre et al., 2016; Parmaksiz and Kim, 2025). Its functional effects depend critically on the precise localization of its receptors, as the spatial distribution of oxytocin receptors determines the neural circuits engaged and the behavioral outcomes elicited. By making brain expression more reliant on distal regulatory sequences, *Oxtr* becomes subject to a wider range of cis- and trans-acting influences, rendering its expression in the brain inherently more variable across individuals and species. This variability allows oxytocin signaling to recruit novel neural circuits and modulate diverse aspects of social behavior, thereby contributing to the evolutionary diversification of sociality. Future studies should aim to resolve the precise mechanisms underlying this variability—for example, by identifying which distal regulatory elements physically interact with *Oxtr* within brain-specific chromatin architectures.

### Ideas and Speculation

In summary, our results revealed that region-specific expression of *Oxtr* in the brain requires long-distance interactions between proximal and distal cis-regulatory elements. In contrast, many other genes remained consistent across BAC-transgenic lines, indicating that proximal regulatory sequences within the BAC were sufficient to drive faithful expression in these tissues, and that this expression was resistant to positional effects.

Our findings support a broader hypothesis: genes vary in their regulatory robustness and evolutionary flexibility. We propose a framework dividing genes into two broad classes based on their transcriptional architecture—rigid and flexible genes. Rigid genes— perhaps comprising the majority of the genome—maintain core life-sustaining functions (e.g., metabolism, motor function, development, reproduction) and are typically regulated by compact proximal elements. Their expression is stable across species, and across individuals within a same species. For such genes, BAC constructs often recapitulate endogenous expression reliably—examples include CamKII, Protein Kinase C Delta(Pkcd), and other genes characterized in large-scale projects like GENSAT.

In contrast, a subset of genes are flexible genes—such as neuromodulatory receptors like *Oxtr* and potentially *Avpr1a* and *Drd4 (Dopamine receptor D4)*— possess inherently more fragile or flexible transcriptional regulation. Their expression is highly sensitive to multiple factors including the chromatin landscape and complex long-range regulatory interactions. We hypothesize that this regulatory sensitivity is not a flaw but an evolutionary feature—what we term transcriptional evolvability. Subtle genetic or epigenetic changes, such as SNPs, structural variants, or chromosomal rearrangements, may lead to variation in neuromodulator receptor expression, thereby altering the modulation of preexisting neural circuits. Such changes could unmask cryptic behavioral phenotypes or give rise to novel social traits without requiring changes to core neural architecture. For example, caregiving circuits might evolve into pair bonding, empathy, or romantic attachment, while defensive or aggressive circuits might become recontextualized into behaviors like jealousy or mate guarding.

This model of transcriptional evolvability helps explain why *Oxtr* and *Avpr1a* expression varies so markedly across and within species, while their ligands (*Oxt* and *Avp*) remain stably expressed. The functional consequence of receptor variability is amplified by the secreted and diffusive nature of these neuromodulators: even small shifts in receptor localization can profoundly alter which neural circuits are engaged and which behaviors are modulated. It also raises the possibility that the capacity for behavioral innovation is, at least in part, encoded in the regulatory fragility of a small set of genes. Our model carries profound implications. It suggests that the evolution of complex social behavior is driven not only by neural circuitry or protein function, but by the regulatory architecture of a small set of flexible genes. These genes serve as behavioral tuning knobs—sensitive to the chromosomal environment, and capable of generating diverse phenotypes in response to evolutionary or environmental pressures. This model also offers insight into variability in human social phenotypes, including those observed in psychiatric conditions such as autism or social anxiety disorders, where subtle changes in receptor gene regulation may lead to major phenotypic shifts. Further research into the regulatory logic of *Oxtr, Avpr1a*, and similar genes may reveal a generalizable mechanism by which genome structure enables behavioral diversity and adaptation across the animal kingdom.

## Materials and Methods

### *pvOxtr-P2A-Cre* BAC vector construction

The vole BAC clone (GeneBank: DP001214.1) containing the *Oxtr* locus was obtained from a prairie vole BAC library (CHORI-232) (McGraw et al., 2010; McGraw et al., 2012). The NLS (nuclear localization signal)-Cre-Frt-Amp-Frt cassette was obtained from Dr. Takuji Iwasato, and was slightly modified to introduce a P2A sequence (Kim et al., 2011). The P2A sequence was introduced using PCR with the following primers: fw5’- ATGGGAAGCGGAGCTACTAACTTCAGCCTGCTGAAGCAGGCTGGAGA CGTGGAGGAGAACCCTGGACCTCTCGAAACTGACAGGAGAACCACC-3’; rv5’-AGACTGGAGTCCGCATAGCCCCCCTCCCCCGCCCCAGGCGCGGTGGGCC AGGCAGGTGGCTCACCTTGACCAAGTTGCTGAAGTTCCTATTCC-3’. The P2A sequence is underlined. And the resulting amplified cassette had the P2A upstream of the NLS-Cre. Then the P2A-NLS-Cre-Frt-Amp-Frt cassette was inserted in-frame right before the stop codon of Oxtr in the BAC clone using the Red/ET Recombineering kit (Gene Bridges GmbH, Heidelberg Germany) after adding homologous arms by PCR with the following primers: fw5’-CACCTTCGTCCTGAGTCGCCGCAGCTCCAGCCAGAGGAGCTG CTCTCAACCATCTTCAGCAATGGGAAGCGGAGCTACTAAC-3’; rv5’-GAGGAGAGGGATACACACCAATAGGCACCTTATACAACTCCACGCAC GGCCACCAGGGGCAGACTGGAGTCCGCATAGCC-3’. The ampicillin selection marker was deleted with the 706-Flpe-induced recombination method (Gene Bridges Heidelberg Germany). A correctly modified BAC clone was verified by using PCR at both the 5’ and 3’ junctions of the targeted insertion with the following primer pair: fr5’-GCCTTCATCATCGCCATGCTCTT-3’(p2) and rv5’-GATGGCTGAGTG ACTGGCATCT-3’(p8); The construct was further verified by sequencing using the following 7 primers specific either for BAC vector or the p2A-NLS-Cre fragment: 5’- CCTGCAGCCAACTGGAGCTTC-3’(p1); 5’- CTTCCTTGGGCGCATTGACGTC-3’(p5); 5’-TACCTGTTTTGCCGGGTCAG-3(p4); 5’-GCCTTCATCATCGCCATGCTCTT-3’(p2); 5’-CTGACCCGGCAAAACAGGTA-3’(p7) ; 5’-TCCGGTTATTCAACTTGCACCATGC-3’(p6); 5’-GATGGCTGAGTGACTGGCATCT-3’(p8). *pvOxtr-P2A-Cre* BAC vector was generated in this study; no RRID is yet available.

### Animals

#### pv*Oxtr-P2A-Cre* BAC transgenic mouse lines (Koi lines)

pv*Oxtr-P2A-Cre* BAC vector was digested with NotI for linearization and to remove the pTARBAC vector backbone. The pv*Oxtr-P2A-Cre* BAC linearized fragment was purified using CL-4B sepharose (Sigma-Aldrich), and injected into pronuclei of C57BL/6J zygotes. Mice carrying the BAC transgene were identified and confirmed using PCR with 5 pairs of primers: fr5’- TCGACCAGGTTCGTTCACTC-3’(p3) + p7; p2+ p6; p2 +p7; p4 +p8; p2+p8. 8 lines (Koi-1∼Koi-8) were confirmed with successful transgene inheritance. These lines were maintained on a congenic C57BL/6J background by backcrossing with wildtype C57BL/6J (RRID: IMSR_JAX:000664) for assessments of BAC DNA copy numbers and initial transgene expression analysis. pv*Oxtr-P2A-Cre* BAC transgenic mouse lines (Koi lines) were generated in this study; no RRID is yet available.

The ***ROSA-26-NLS-LacZ* mouse** line (Zhang et al., 2016) was obtained from Itohara Lab at RIKEN (RRID:SCR_001065); no RRID is assigned to this line.

The ***mOxtr-Ires-Cre* knock-in** (Hidema et al., 2016) and ***mOxtr^−/−^* mutant** (Takayanagi et al., 2005) lines were obtained from Dr. Katsuhiko Nishimori at Tohoku University, and both were backcrossed with C57BL/6J for more than 10 generations. Each Koi line and *mOxtr-Ires-Cre* knock-in mice were crossed with *ROSA-26-NLS-LacZ* mice to generate Cre^+^, LacZ^+^ double positive mice for further gene expression analysis. Female Koi mice were crossed with male m*Oxtr*^−/−^ to produce offspring with the genotype of pv*Oxtr-P2A-Cre*(Koi): m*Oxtr*^+/−^. These offspring were then bred to produce Koi: m*Oxtr*^−/−^ mice expressing pv*Oxtr* while lacking endogenous *mOxtr*, and these mice were used for autoradiography, qRT-PCR and behavioral tests. All mice were generated using continuously housed breeder pairs and P21 as the standard weaning date. All the animal procedures were approved by University of Tsukuba Animal Care and Use Committee. Mice were housed under constant temperature and light condition (12 h light and 12 h dark cycle), and received food and water ad libitum.

### Determination of transgene copy number

Custom Taqman® MGB probes were synthesized by oligoJp (Thermo Fisher, Japan) for detecting the transgenic *Cre* and the mouse *Jun* gene (internal control). The following primer pairs and probes were used: for the *Cre* assay, fr5’- ATGACTGGGCACAACAGACAAT-3’; rv5’- CGCTGACAGCCGGAACAC-3’; probe: 5’-FAM-AAACATGCTTCATCGTCGGTC CGG-MGB-3’; for *Jun* assay, fr5’- GAGTGCTAGCGGAGTCTTAACC-3’; rv5’-CTCCAGACGGCAGTGCTT-3’; probe: 5’-VIC-CTGAGCCCTCCTCCCC-MGB-3’. Real-time PCR was performed using an Applied Biosystem7500. Transgene copy number was determined through absolute quantification (standard curve method) following the protocol described in (Chandler et al., 2007). Briefly, 2 microliters (20 ng) of genomic DNA samples or copy number standards were analyzed in a 20μL reaction volume with two primer-probe sets (*Cre, Jun*). The data from transgenic samples were then compared to a standard curve of calibrator samples that are generated by diluting purified BAC DNA (linearized) over a range of known concentrations into wild-type mouse genomic DNA. In addition, no-template controls were included in each experiment. All reactions were performed in triplicate.

### Histology

Adult mice (3-4month old), 4 male and 4 female from each line/ generation were analyzed for 3 generations. Mice were perfused intracardially with 4% formalin in 0.1M sodium phosphate buffer. Brains were embedded in 2% agarose in 0.1M PB and cut into 100-m-thick sections with a Micro-slicer (Dosaka, Kyoto, Japan). Mammary glands were isolated from virgin female mice (2month old), 2 mice/each line were analyzed. The brain slices and whole mount mammary glands preparation were stained in X-gal solution (5 mM K3FeCN6, 5mMK4FeCN6,2mMMgCl2, 0.02% NP-40, 0.01% Na-deoxycholate, 1 mg/ml X-gal in 0.1M PB) at 37°C for 6 hours and then were stained with hematoxylin. All experiments were done with positive and negative controls (Cre^+^, LacZ^−^) to monitor the reliability of the X-gal staining. Images were caught by Nanozoomer 2.0-HT slide scanner (Hamamatsu Photonics).

### Receptor Autoradiography

OXTR autoradiography was performed as previously described (Inoue et al., 2022). Briefly, freshly frozen brains were stored at −80°C. Coronal sections were cut in a cryostat and 20 µm sections were collected and then stored at −80°C until use in autoradiography. Brain sections were removed from −80°C storage and air dried, and then fixed for two minutes with 0.1% paraformaldehyde in PBS at room temperature, and rinsed twice in 50 mM Tris buffer, pH 7.4, to remove endogenous OT. They were then incubated in 50 pM 125I-OVTA (2200 Ci/mmol; PerkinElmer; Boston, MA), a selective, radiolabeled OXTR ligand, for one hour. Unbound 125I-OVTA was then washed away with Tris-MgCl_2_ buffer (50 mM Tris plus 2% MgCl2, pH 7.4) and sections were air dried. Sections were exposed to BioMax MR film (Kodak; Rochester, New York) for five days. Digital images were obtained with a light box and a Cannon camera (Cannon 6D MarkII, Japan). The brightness and contrast of representative images were equally adjusted for all autoradiography images within a panel using Adobe Photoshop (RRID:SCR_014199).

### Fluorescent RNAscope *in situ* hybridization for *Oxtr* mRNA localization

RNAscope Multiplex Fluorescent Reagent Kit v2 (Advanced Cell Diagnostics, Newark, California; Cat. No. 323270) was used according to the manufacturer’s instructions. Adjacent sections to those utilized for I-125 OVTA receptor binding autoradiography were used. Briefly, sections were thawed and fixed in 10% neutral buffered formalin for 15 minutes followed by treatment with hydrogen peroxide for 10 minutes and RNAscope Protease IV for 30 minutes. Probes specific to prairie vole *Oxtr* mRNA (Cat. No. 500721) were then applied and allowed to hybridize for two hours at 40 °C. After three amplification steps, hybridized probes were labelled with horseradish peroxidase (HRP), and TSA Vivid Dye 570 fluorophore. Sections were counterstained with DAPI for nuclei labeling and coverslipped with ProLong Gold Antifade Mountant (ThermoFisher Scientific; Waltham, Massachusetts). Negative controls were performed using a bacterial dapB probe (Cat. No. 310043), which did not exhibit any nonspecific labeling. Fluorescent RNAscope images were acquired using a Keyence BZ-X710 microscope. Whole sections were captured with 10x objective lens in two channels (red and blue), and images were stitched together using Keyence BZX Analyzer Software. Brightness and contrast of the images were adjusted using Photoshop (RRID:SCR_014199).

### Quantification of *Oxtr* mRNA expression by qRT-PCR

Real-time quantitative polymerase chain reaction (RT-qPCR) using an Applied Biosystems 7500 Real-Time PCR system was used to quantify *Oxtr* mRNA expression levels in different brain regions of Koi lines. Total RNA of different brain regions and mammary glands was isolated using the RNeasy extraction kit (Qiagen, Germantown, US), and cDNA was synthesized after Deoxyribonuclease I (Invitrogen, Waltham, US) treatment by using EvoScript Universal cDNA Master (Roche, Penzberg, Germany), and then quantitative RT-PCR was performed with FastStart Universal SYBR Green Master (Roche, Penzberg, Germany). The relative standard curve method was used to obtain the relative quantities of *Oxtr* expression following the manual of Applied Biosystems. To allow the comparation between Koi lines and WT mice, the following primer pair (which can detect the expression of both *mOxtr* and pv*Oxtr*) was used: fr5’- GCCTTTCTTCTTCGTGCAGATG-3’ and rv5′-ATGTAGATCCAGGGGTTGCAG-3′ In addition, to specifically detect the expression of pv*Oxtr*, the primer pair fr5′- GCCTTTCTTCTTCGTGCAGATG-3 and rv5’-AAAGAGGTGGCCCGTGAAC-3′ was used in the 2^nd^ qRT-PCR experiment. GAPDH was used as the endogenous control for both experiments and was amplified by the following primer pair: fr5’- GGGTTCCTATAAATACGGACTGC-3’ and rv5’- CCATTTTGTCTACGGGACGA-3’. Samples were analyzed in triplicates. A non-template control was performed to ensure that there was no amplification of genomic DNA. All of the experimenters were blind to the genotype of the subjects.

### Partner preference test (PPT)

Subjects were housed with 4 age-matched and same-sex littermates until testing at adulthood (2-5 months old). 12-14 homozygous mice were tested for each group. Ovariectomized females were assigned to pair and cohouse with sexually experienced adult wildtype males for 21 days before PPT test using simple randomization. Ovariectomy surgery was performed according to the step-by-step protocol available at STAR Protocols (https://star-protocols.cell.com/protocols/3330). The female was injected with estradiol benzoate (10 μg and 5 μg at 48 h and 24 h before induced mating) and progesterone (500 μg at 4–7 h before induced mating) to ensure high sexual receptivity before mating. Mating was induced on Day 2,9 and16. There was attrition in the study: one Koi-3: mOxtr^−/−^ mouse did not survive the co-housing stage and was therefore excluded from the study.PPT data was analyzed using TimeSSI1 for social interaction test system (O’ Hara & Co., Ltd.). Briefly, the experimental subject was placed in a chamber, in which two pen-boxes (diameter 8cm, height10cm) were located at two diagonally opposite corners. The experimental animal was free to move throughout the chamber and the time spent in close proximity to each pen-box (stay time) is recorded using an automated mouse tracking system (Fig.4 A, B). In the habituation phase, the experimental subject was allowed to explore the chamber for 5 minutes with both pen-boxes empty. In the recognition phase, the partner stimulus animal was restricted in one pen-box, and a novel “stranger” stimulus animal was restricted in another pen-box, and the experimental subject was allowed to explore the chamber for 10 minutes. In the preference phase, the positions of the partner stimulus animal and the “stranger” stimulus animal were switched, and the experimental subject was allowed to explore the chamber for 20 minutes. To assure unbiased design, pen-box assignments were counterbalanced for the diagonal positions. All of the experimenters were blind to the genotype of the subjects.

### Parental behavior test

To examine maternal behavior, we performed pup retrieval test with virgin female mice (2- to 4-months old, 15-18mice/line), following the method described in (Stolzenberg et al., 2012) with some modifications. Briefly, 5 days prior to the behavioral testing, each subject was isolated and housed in separate home cages in the test room. Nesting material (about 1.5 g of cotton wool) was provided for nest making. The tests were conducted in the dark phase of light/dark cycle (12 hours/12 hours). On the day of the test, each subject was placed in the recording area and allowed 15 minutes for habituation. The stimulus pups (age: P2-P3) were collected from a group of donor mothers immediately before the start of the experiment. A retrieval test began with the placement of 3 stimulus pups on the side farthest form the nest, and the female’s behavior was recorded by an ARNAN 4 channel security system for 15 minutes. Pup retrieval was defined as picking up the pup and bringing it to the nest. If the subject did not complete the retrieval for 3 pups in 15 minutes, the video recording time was extended to 30 minutes. The latency to retrieve each pup (1st, 2nd, 3rd) and the time spent crouching on the pups were time-stamped and calculated manually. A latency of 1800 seconds was assigned if the pup-retrieval was not completed in 30 minutes. Crouching was defined as the mouse supporting itself in a lactation-position over the pups in the nest. Crouching time on a single pup, two pups, and three pups were summed up as total crouching time. All of the experimenters were blind to the genotype of the subjects.

### 3D chromatin verification on Hi-C datasets of humans and mice

The 3D genome structure tracks were obtained from 4DNucleome Consortium (https://www.4dnucleome.org/) where datasets were systematically reanalyzed following recommended standard protocols. The datasets IDs and sample information are provided in the supplementary material (Supplementary Table1). 3D Genome Browser(Wang et al., 2018) and WashU Epigenome Browser(Li et al., 2022) were employed for visualizations.

### Experimental design and statistical analysis

The sample sizes were determined by G*Power 3 program (Faul et al., 2007) (RRID:SCR_013726) based on pilot experiments. They are also within the range that is generally accepted in the field. All experiments were performed and analyzed blind to the genotypes and no data point was excluded. All reported sample numbers (n) represent independent biological replicates that are the numbers of tested mice or tissue. All the statistical analyses were performed by SPSS21(IBM). Mixed ANOVA was used for analyzing PPT data and pup retrieval data. Mauchly’s test of sphericity was used to test whether or not the assumption of sphericity was met in repeated measures. Greenhouse-Geisser correction was applied when the sphericity was violated. Two-way ANOVA was used for analyzing qRT-PCR region specific data across different mouse lines. One-way ANOVA was used for analyzing crouching time in the parental behavioral test. If there was a significant main effect of an independent factor, post hoc test was used to do multiple comparisons. If there was a significant interaction between within-subject factor and between-subject factor, post hoc pairwise comparisons were applied to analyze the simple main effects. Bonferroni correction was used for all the post hoc tests.

## Acknowledgements

This work was supported by the RIKEN Incentive Research Project (100226201701100443) to Q.Z., the Brain Science Project, Center for novel science initiatives, National institutes of natural sciences (BS291003) to Q.Z., the RIKEN Aging Project (10026-201701100263-340120) to Q.Z., the JSPS Kakenhi Grant-in-Aid for Young Scientists (17K18362) to Q.Z., the JSPS Kakenhi Grant-in-Aid for Challenging Research (19K21807) to Q.Z., the JSPS Kakenhi Grant-in-Aid for Challenging Research (22K19478)to Q.Z., the JSPS Kakenhi Grant-in-Aid for Scientific Research (25K10198) to Q.Z., the Takeda Medical Research Grant to Q.Z., the International Education and Research Laboratory Program of University of Tsukuba to L.Y. and Q.Z.. We thank Dr. Katsuhiko Nishimori (Tohoku University and Fukushima Medical University, Japan) for providing *mOxtr*-Ires-Cre knock-in line and *mOxtr*−/− mutant line; Dr. Cary Lai (Indiana University, USA), Dr. Takuji Iwasato(National Institute of Genetics, Japan) and Dr. Seiya Mizuno(University of Tsukuba, Japan) for their advice on BAC vector construction; Dr. Volkhard Maeckel (CERN, Switzerland) for helping with image capturing; Dr. Yong Zhang (Tongji University, China) for the stimulating discussion about 3D chromatin structure, and we thank the Laboratory of Animal Resource Center of University of Tsukuba for pronuclear injection.

## Author Contributions

Q.Z. and L.Y. conceived and designed the study. Q.Z., M.T., L.N., K.I., L.K., M.P., Y.N., M.S., and T.S. performed experiments. Q.Z., M.T., L.N., K.I., L.K., M.P., K.N. and L.Y analyzed data. S.I. provided guidance for transgene vector construction. Q.Z. and L.Y. supervised the project and wrote the manuscript. All authors contributed to discussion and approved the final version.

## Supplementary Information

**Supplementary Fig.1.**
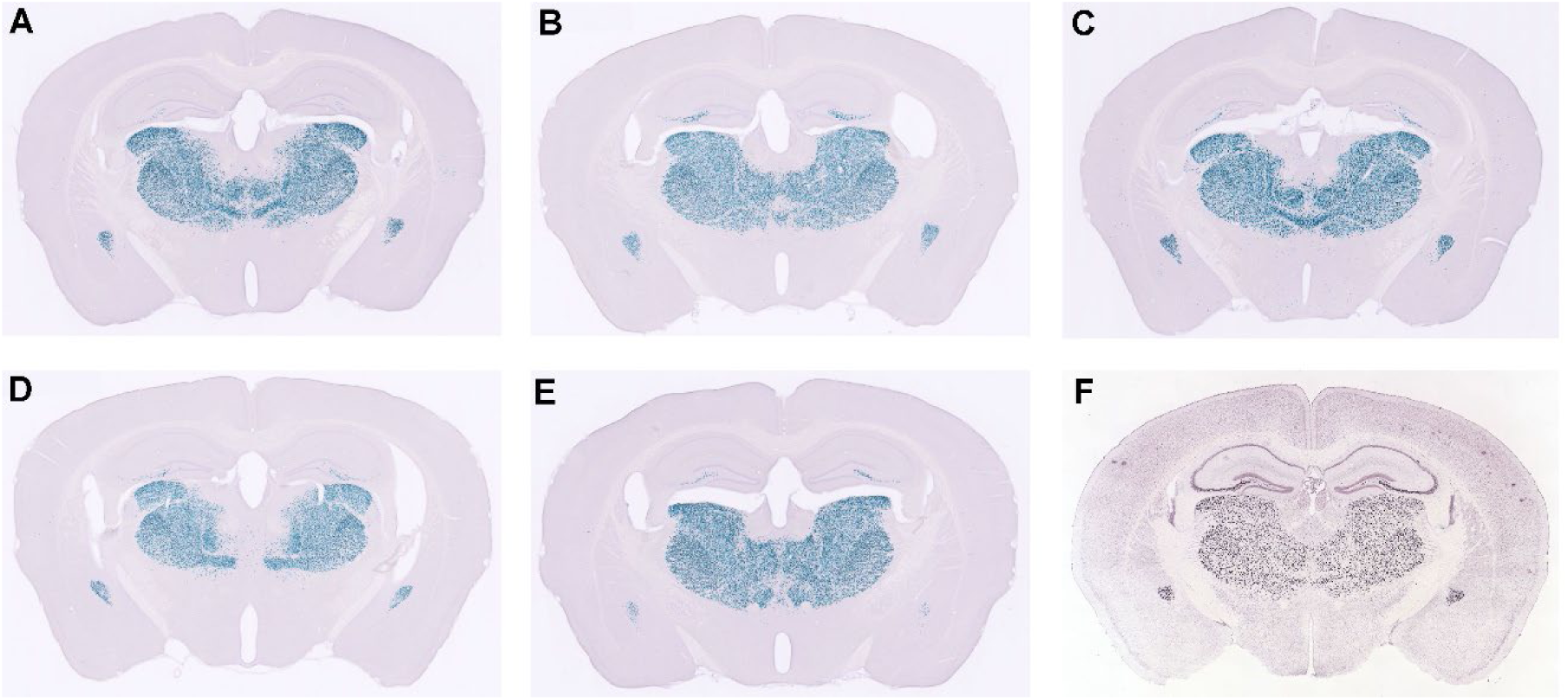
Expression pattern of the *Pkcd-Cre* BAC transgene. All five independent *Pkcd-Cre* BAC transgenic mouse lines (A–E) exhibited expression patterns that closely resembled the endogenous *Pkcd* expression, as shown by in situ hybridization (F). The *Pkcd-Cre* BAC construct was generated using the same strategy as that employed for the Koi lines, incorporating the entire intergenic regions both upstream and downstream of the *Pkcd* gene.

**Supplementary Fig.2.**
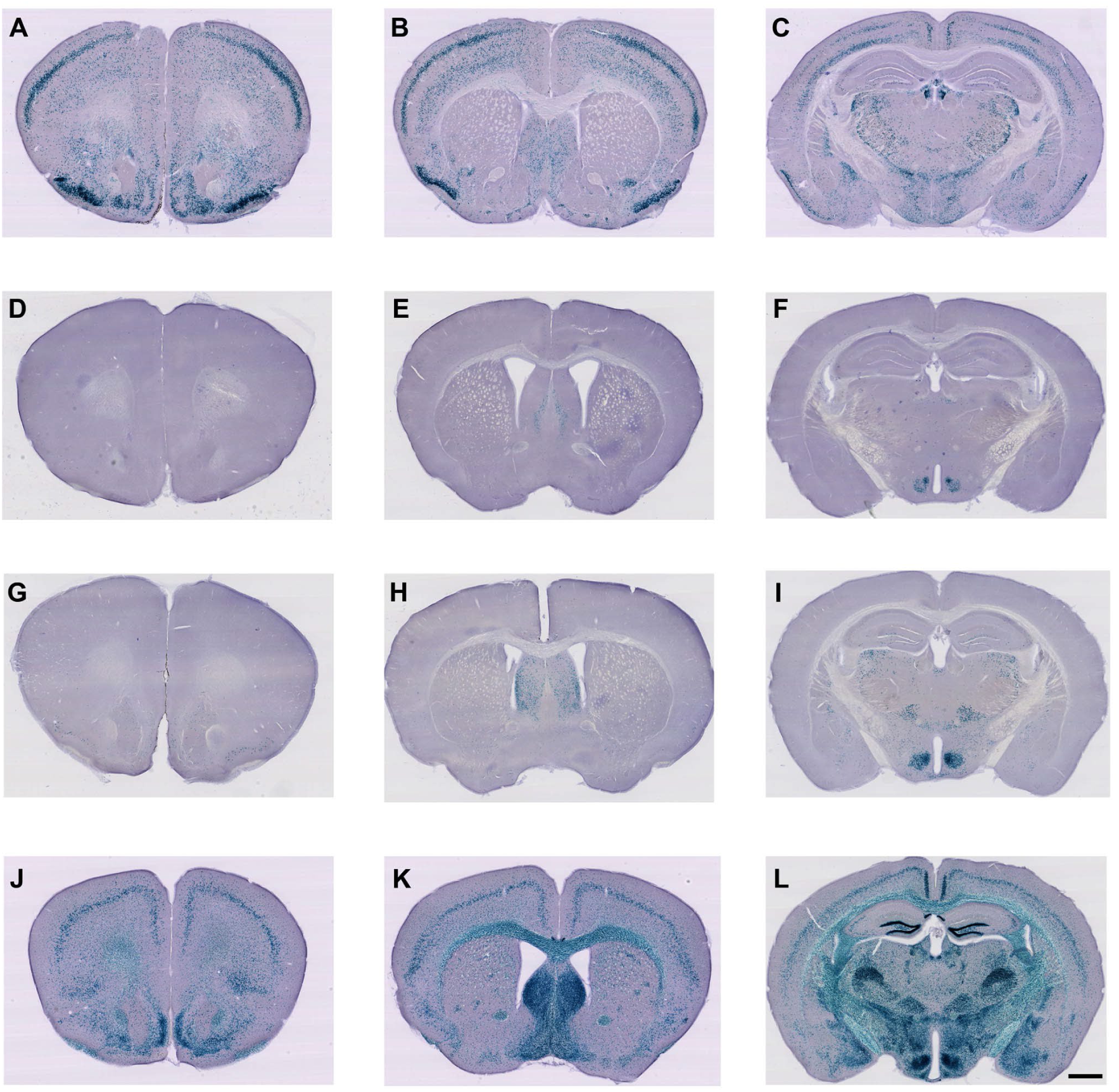
Transgenic CRE mediated Lac-Z reporter gene expression in the brain of Koi lines. X-gal staining of the brains from the double positive (*Cre*+, *LacZ*+) offspring of Koi-5 (A-C), Koi-6 (D-F), Koi-7 (G-I), Koi-8 (J-L). Scale bar=1mm

**Supplementary Fig.3.**
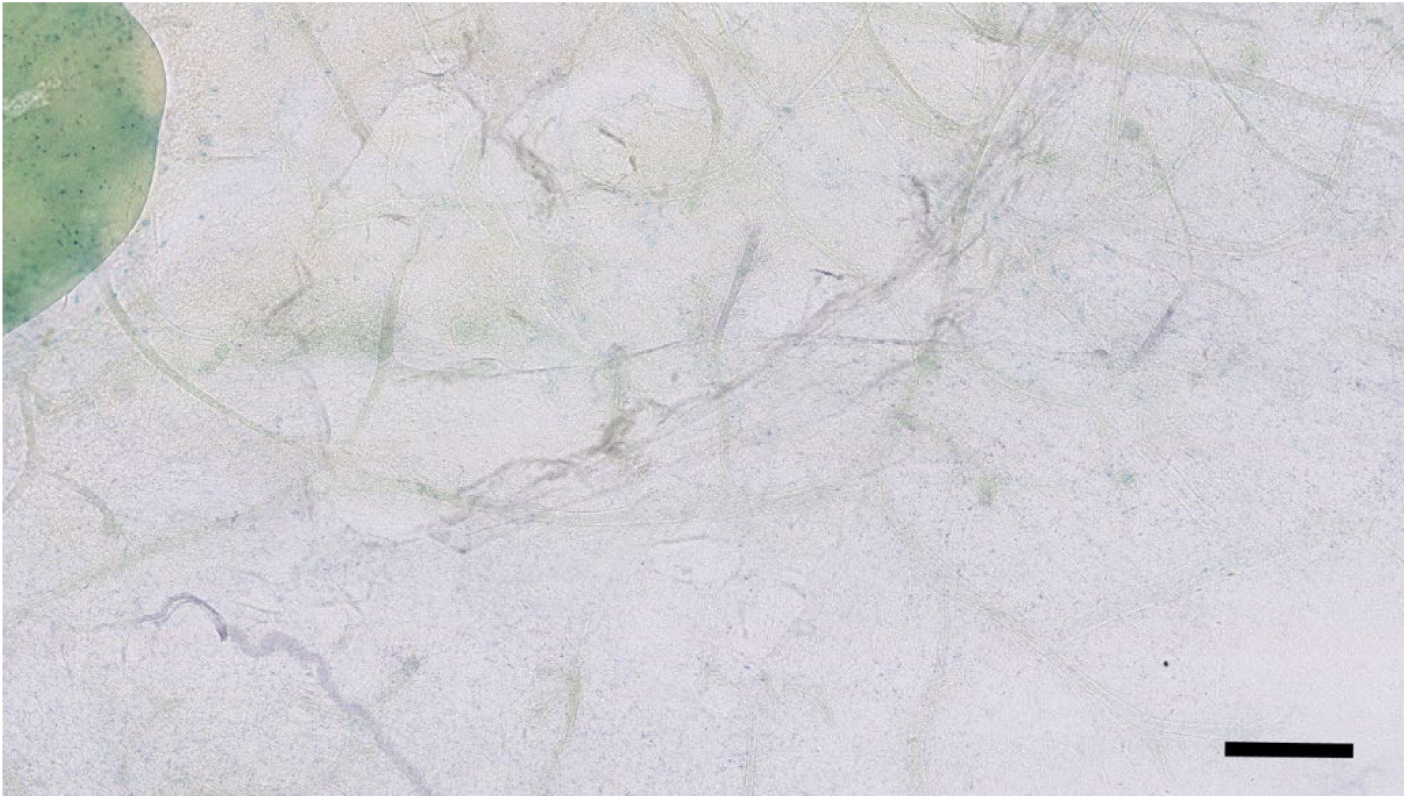
Negative control of whole-mount mammary gland staining. X-gal staining of a mammary gland from (*Cre*-, *LacZ*+) offspring from heterozygous Koi-4 was shown. The lymph node showed some blue signal, indicating that the lymph node could be stained nonspecifically. All the epithelium ducts did not show any blue signal, verifying the specificity of duct labeling of X-gal staining. Scale=500um

**Supplementary Fig.4.**
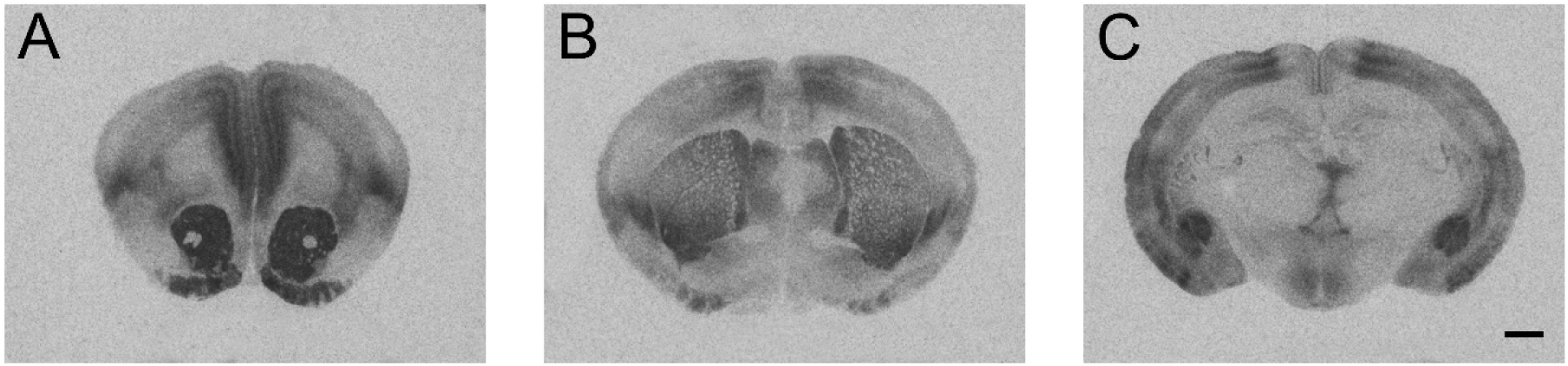
OXTR expression in the prairie vole brain. Autoradiographs illustrate the distribution of OXTR binding in the adult prairie vole brain at (A)prefrontal cortex level, (B) septum and striatum level, and (C) Basolateral amygdala level. Scale bar=1mm. These autoradiographs were developed using BioMax MR film, which had since been discontinued. The difference in image appearance compared with the transgenic mouse autoradiographs shown in Fig3. Reflects the use of different film types. The cross-species comparison of OXTR expression was intended as qualitative assessments of spatial distribution rather than direct quantitative comparisons.

**Supplementary Fig.5.**
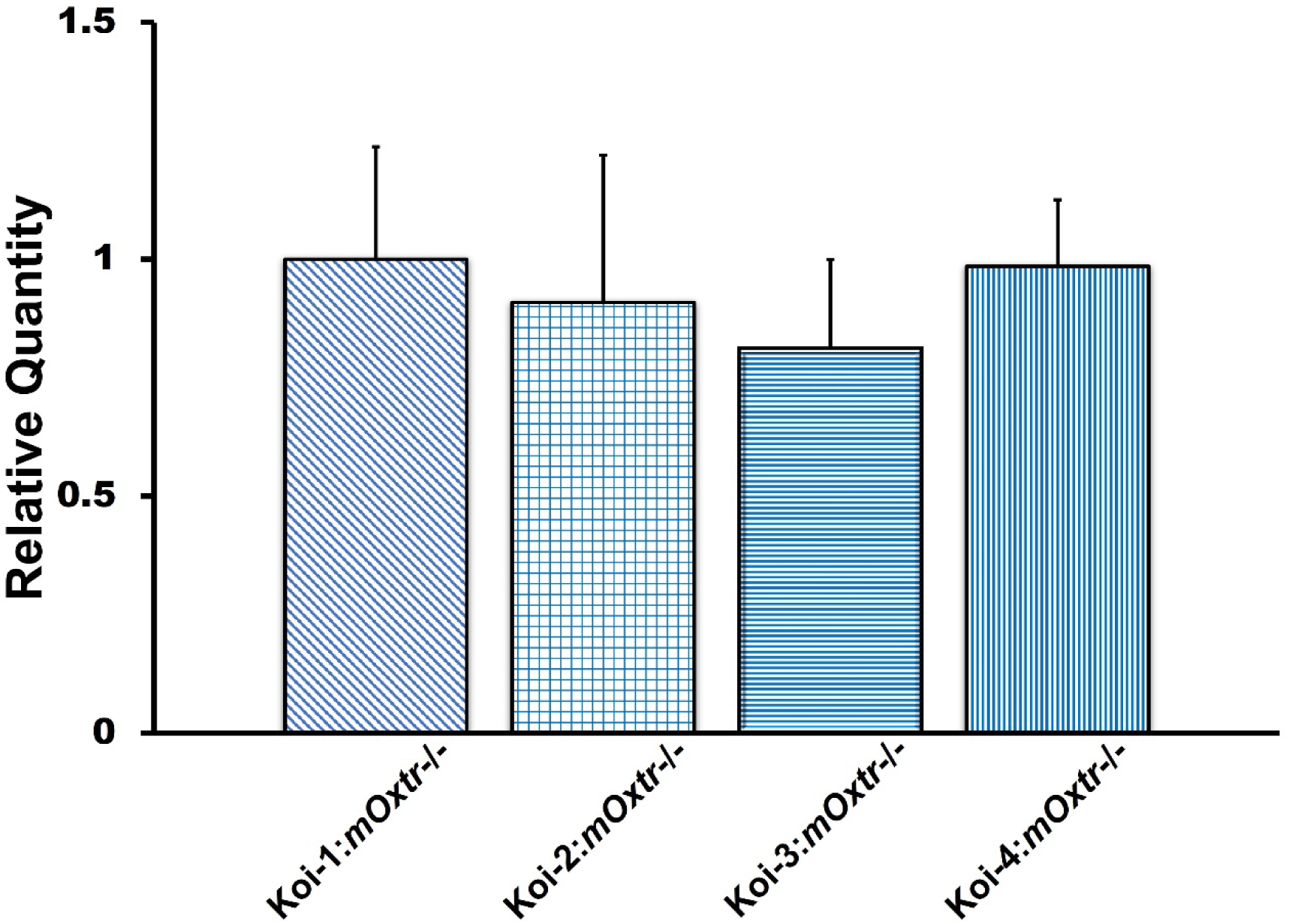
Quantitative PCR analysis of *Oxtr* mRNA expression in mammary gland of Koi lines. A primer set specifically recognizing pv*Oxtr.* was used to analyze *Oxtr* mRNA in Koi-1: m*Oxtr*^−/−^, Koi-2: m*Oxtr*^−/−^, Koi-3: m*Oxtr*^−/−^ and Koi-4: m*Oxtr*^−/−^ mic, n=6 for each genotype. There was no significant difference in Oxtr expression among the Koi lines (one-way ANOVA, F (3, 20) = 0.307, p = 0.82; Supplementary Fig. 5)

**Supplementary Fig.6.**
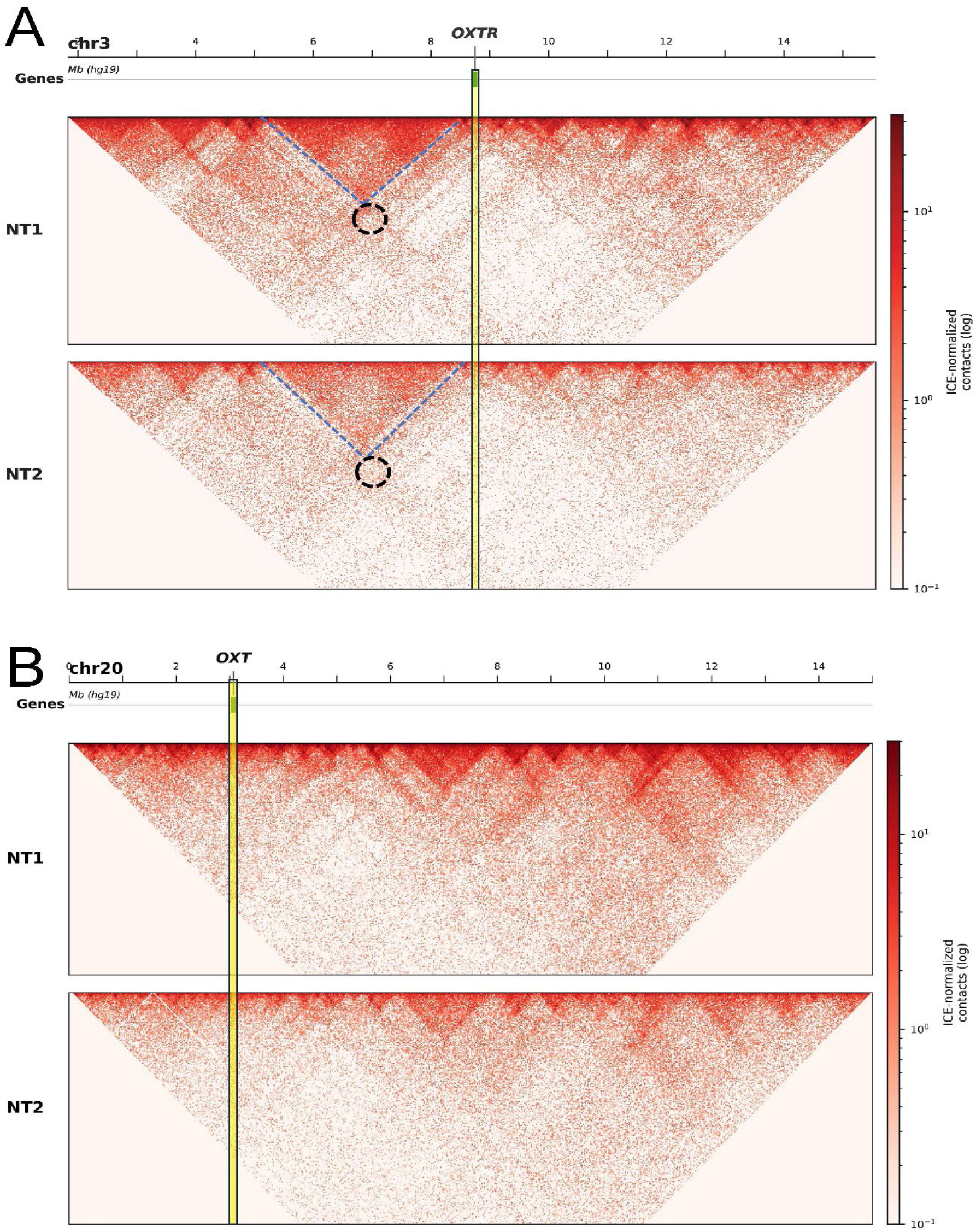
Heatmap of the chromatin contacts surrounding *Oxtr* and *Oxt* in human mammary tissue. Heatmaps show chromatin contacts surrounding *OXTR* and *OXT* in normal human breast tissue from two donors (NT1 and NT2) at megabase-scale resolution. The *OXTR* locus is located at a prominent TAD boundary (A), whereas the *OXT* region lacks a clear TAD structure (B). TAD structures surrounding *OXTR* are indicated by blue dashed lines. No prominent long-range chromatin interactions are observed surrounding *OXTR*; the signal within the black-circled region is comparable to the background signal in the heatmap, which matches the mouse hepatocyte/sperm panels, but not the brain tissue panels in Fig.7A. These features were consistently identified across both donors using independent insulation-based and TopDom analyses. The human breast Hi-C dataset has substantially lower sequencing depth than the mouse Hi-C datasets used in Fig. 7, resulting in lower-resolution heatmaps.

### Supplementary Methods

#### Hi-C data analysis for human breast tissue (Supplementary Fig.6)

Because no publicly available Hi-C dataset from mouse mammary gland was identified in the 4DNucleome, ENCODE, or GEO repositories, we used Hi-C data from normal human breast tissue for Supplementary Fig. 6 (Choppavarapu et al., 2025). Hi-C data from two normal breast tissue donors (NT1 and NT2) were obtained through direct communication with the authors, who kindly provided access to the data from their repository (github.com/KunFang93/TumorSpecific3DChromatinDomain). The data consisted of HiC-Pro-processed, ICE-normalized contact matrices at 20-kb resolution (hg19). Only normal tissue samples were analyzed; tumor samples from the same dataset were excluded. Because the deposited matrices did not include the corresponding bin-coordinate file, the hg19 20-kb bin map was reconstructed from the reference chromosome sizes and validated against the dimensions of the contact matrices (154,795 bins genome-wide). The matrices were then loaded into cooler (v0.10.4) for visualization. The genomic positions of *OXTR* (chr3:8,811,300) and *OXT* (chr20:3,052,265) were independently confirmed using the UCSC RefSeq annotation for hg19. Genome-wide insulation scores were independently calculated using cooltools with a 500-kb window to further evaluate the local domain organization surrounding *OXTR* and *OXT*.

**Supplementary Table.1.** The datasets IDs and sample information for mouse 3D chromatin verification.

|  | track | ID | Cell/tissue | Organism | Restr. Enz. | Database | Database ID | Browser | Publication |
| --- | --- | --- | --- | --- | --- | --- | --- | --- | --- |
| Hi-C | 1 | Whole brain | whole brain | mm10 | DNaseI | 4DN | 4DNFIQ48NYOW | WashU | Deng 2015 |
|  | https://4dn-open-data-public.s3.amazonaws.com/fourfront-webprod/wfoutput/60d98cc6-4583-4ebe-9c5a-77fc421ef560/4DNFIQ48NYOW.hic |  |  |  |  |  |  |  |  |
|  | 2 | NPC | NPC | mm10 | DpnII | 4DN | 4DNFIHW8NTQX | WashU | Bonev 2017 |
|  | https://4dn-open-data-public.s3.amazonaws.com/fourfront-webprod/wfoutput/219497b5-3f35-473b-9d4e-e1cf21c69561/4DNFIHW8NTQX.hic |  |  |  |  |  |  |  |  |
|  | 3 | Cortex | cortex | mm10 | Mbol | 4DN | 4DNFII3JV8I1 | WashU | Du 2017 |
|  | https://4dn-open-data-public.s3.amazonaws.com/fourfront-webprod/wfoutput/268b7d52-9655-474c-9467-8ba31bb2195c/4DNFII3JV8I1.hic |  |  |  |  |  |  |  |  |
|  | 4 | ESC | ESC | mm10 | DpnII | 4DN | 4DNFIU8AF5ZY | WashU | Bonev 2017 |
|  | https://4dn-open-data-public.s3.amazonaws.com/fourfront-webprod/wfoutput/c2eaf9bf-9584-4cec-8685-bd74038a6c01/4DNFIU8AF5ZY.hic |  |  |  |  |  |  |  |  |
|  | 5 | Hepatocyte | hepatocyte | mm10 | HindIII | 4DN | 4DNFIFULDMGN | WashU | Schwarzer 2017 |
|  | https://4dn-open-data-public.s3.amazonaws.com/fourfront-webprod/wfoutput/57ecaa79-5c9c-49c7-b841-0a4bf3117555/4DNFIFULDMGN.hic |  |  |  |  |  |  |  |  |
|  | 6 | Sperm | sperm | mm10 | Mbol | 4DN | 4DNFIN8F14CS | WashU | Du 2017 |
|  | https://4dn-open-data-public.s3.amazonaws.com/fourfront-webprod/wfoutput/27f54fcb-54fe-41a4-b25a-2f8944c89044/4DNFIN8F14CS.hic |  |  |  |  |  |  |  |  |
|  | Fig | Regions | Genes |  |  |  |  |  |  |
| WashU | A | chr6:104149809-119528368 | Oxtr |  |  |  |  |  |  |
|  | B | chr2:121804885-139358632 | Oxt |  |  |  |  |  |  |

The database source of each mouse sample, analysis and visualization tool information, and the region analyzed was listed.

## Data and Materials Sharing

Original data created for the study are or will be available in a persistent repository upon publication Repository Name and DOI/accession number(s):

**zhang, qi, “Dataset - Hi-c analysis for Oxtr and Oxt”, Mendeley Data, V1, doi: 10.17632/jdn6z5m6zs.1**

**zhang, qi, “Dataset for the analysis of koi mice”, Mendeley Data, V1, doi: 10.17632/fjztcwtj2m.1**

All data is included in the manuscript and/or supporting information

## References

Ahern TH, Olsen S, Tudino R, Beery AK (2021) Natural variation in the oxytocin receptor gene and rearing interact to influence reproductive and nonreproductive social behavior and receptor binding. Psychoneuroendocrinology 128:105209.

Amadei EA, Johnson ZV, Jun Kwon Y, Shpiner AC, Saravanan V, Mays WD, Ryan SJ, Walum H, Rainnie DG, Young LJ, Liu RC (2017) Dynamic corticostriatal activity biases social bonding in monogamous female prairie voles. Nature 546:297–301.

Barrett CE, Arambula SE, Young LJ (2015) The oxytocin system promotes resilience to the effects of neonatal isolation on adult social attachment in female prairie voles. Transl Psychiatry 5:e606.

Beagan JA, Phillips-Cremins JE (2020) On the existence and functionality of topologically associating domains. Nature Genetics 52:8–16.

Beil J, Fairbairn L, Pelczar P, Buch T (2012) Is BAC transgenesis obsolete? State of the art in the era of designer nucleases. J Biomed Biotechnol 2012:308414.

Berendzen KM, Sharma R, Mandujano MA, Wei Y, Rogers FD, Simmons TC, Seelke AMH, Bond JM, Larios R, Goodwin NL, Sherman M, Parthasarthy S, Espineda I, Knoedler JR, Beery A, Bales KL, Shah NM, Manoli DS (2023) Oxytocin receptor is not required for social attachment in prairie voles. Neuron 111:787–796.e784.

Bian Q, Belmont AS (2010) BAC TG-EMBED: one-step method for high-level, copy-number-dependent, position-independent transgene expression. Nucleic Acids Research 38:e127–e127.

Carcea I et al. (2021) Oxytocin neurons enable social transmission of maternal behaviour. Nature 596:553–557.

Chandler KJ, Chandler RL, Broeckelmann EM, Hou Y, Southard-Smith EM, Mortlock DP (2007) Relevance of BAC transgene copy number in mice: transgene copy number variation across multiple transgenic lines and correlations with transgene integrity and expression. Mamm Genome 18:693–708.

Chen Z, Han Y, Ma Z, Wang X, Xu S, Tang Y, Vyssotski AL, Si B, Zhan Y (2024) A prefrontal-thalamic circuit encodes social information for social recognition. Nature Communications 15:1036.

Choppavarapu L, Fang K, Liu T, Ohihoin AG, Jin VX (2025) Hi-C profiling in tissues reveals 3D chromatin-regulated breast tumor heterogeneity informing a looping-mediated therapeutic avenue. Cell Reports 44.

Crook ZR, Housman D (2011) Huntington’s disease: can mice lead the way to treatment? Neuron 69:423–435.

Dekker J, Rippe K, Dekker M, Kleckner N (2002) Capturing chromosome conformation. Science 295:1306–1311.

Dixon JR, Selvaraj S, Yue F, Kim A, Li Y, Shen Y, Hu M, Liu JS, Ren B (2012) Topological domains in mammalian genomes identified by analysis of chromatin interactions. Nature 485:376–380.

Faul F, Erdfelder E, Lang AG, Buchner A (2007) G*Power 3: a flexible statistical power analysis program for the social, behavioral, and biomedical sciences. Behav Res Methods 39:175–191.

Ferrante RJ (2009) Mouse models of Huntington’s disease and methodological considerations for therapeutic trials. Biochimica et Biophysica Acta (BBA) - Molecular Basis of Disease 1792:506–520.

Froemke RC, Young LJ (2021) Oxytocin, Neural Plasticity, and Social Behavior. Annu Rev Neurosci 44:359–381.

Gerfen CR, Paletzki R, Heintz N (2013) GENSAT BAC cre-recombinase driver lines to study the functional organization of cerebral cortical and basal ganglia circuits. Neuron 80:1368–1383.

Gibcus Johan H, Dekker J (2013) The Hierarchy of the 3D Genome. Molecular Cell 49:773–782.

Gilligan P, Brenner S, Venkatesh B (2003) Neurone-specific expression and regulation of the pufferfish isotocin and vasotocin genes in transgenic mice. J Neuroendocrinol 15:1027–1036.

Gong S, Doughty M, Harbaugh CR, Cummins A, Hatten ME, Heintz N, Gerfen CR (2007) Targeting Cre recombinase to specific neuron populations with bacterial artificial chromosome constructs. J Neurosci 27:9817–9823.

Gong S, Zheng C, Doughty ML, Losos K, Didkovsky N, Schambra UB, Nowak NJ, Joyner A, Leblanc G, Hatten ME, Heintz N (2003) A gene expression atlas of the central nervous system based on bacterial artificial chromosomes. Nature 425:917–925.

Grinevich V, Knobloch-Bollmann HS, Eliava M, Busnelli M, Chini B (2016) Assembling the Puzzle: Pathways of Oxytocin Signaling in the Brain. Biol Psychiatry 79:155–164.

Heintz N (2004) Gene Expression Nervous System Atlas (GENSAT). Nature Neuroscience 7:483–483.

Hidema S, Fukuda T, Hiraoka Y, Mizukami H, Hayashi R, Otsuka A, Suzuki S, Miyazaki S, Nishimori K (2016) Generation of Oxtr cDNA(HA)-Ires-Cre Mice for Gene Expression in an Oxytocin Receptor Specific Manner. J Cell Biochem 117:1099–1111.

Inoue K, Ford CL, Horie K, Young LJ (2022) Oxytocin receptors are widely distributed in the prairie vole (Microtus ochrogaster) brain: Relation to social behavior, genetic polymorphisms, and the dopamine system. Journal of Comparative Neurology 530:2881–2900.

Jurek B, Neumann ID (2018) The Oxytocin Receptor: From Intracellular Signaling to Behavior. Physiol Rev 98:1805–1908.

Kandasamy LC, Tsukamoto M, Banov V, Tsetsegee S, Nagasawa Y, Kato M, Matsumoto N, Takeda J, Itohara S, Ogawa S, Young LJ, Zhang Q (2021) Limb-clasping, cognitive deficit and increased vulnerability to kainic acid-induced seizures in neuronal glycosylphosphatidylinositol deficiency mouse models. Human Molecular Genetics 30:758–770.

Kappel JM, Förster D, Slangewal K, Shainer I, Svara F, Donovan JC, Sherman S, Januszewski M, Baier H, Larsch J (2022) Visual recognition of social signals by a tectothalamic neural circuit. Nature 608:146–152.

Kim JH, Lee SR, Li LH, Park HJ, Park JH, Lee KY, Kim MK, Shin BA, Choi SY (2011) High cleavage efficiency of a 2A peptide derived from porcine teschovirus-1 in human cell lines, zebrafish and mice. PLoS One 6:e18556.

King LB, Walum H, Inoue K, Eyrich NW, Young LJ (2016) Variation in the Oxytocin Receptor Gene Predicts Brain Region-Specific Expression and Social Attachment. Biol Psychiatry 80:160–169.

Leng G, Ludwig M (2008) Neurotransmitters and peptides: whispered secrets and public announcements. J Physiol 586:5625–5632.

Li D, Purushotham D, Harrison JK, Hsu S, Zhuo X, Fan C, Liu S, Xu V, Chen S, Xu J, Ouyang S, Wu AS, Wang T (2022) WashU Epigenome Browser update 2022. Nucleic Acids Research 50:W774–W781.

Lieberman-Aiden E, van Berkum NL, Williams L, Imakaev M, Ragoczy T, Telling A, Amit I, Lajoie BR, Sabo PJ, Dorschner MO, Sandstrom R, Bernstein B, Bender MA, Groudine M, Gnirke A, Stamatoyannopoulos J, Mirny LA, Lander ES, Dekker J (2009) Comprehensive Mapping of Long-Range Interactions Reveals Folding Principles of the Human Genome. Science 326:289–293.

Mack KL, Campbell P, Nachman MW (2016) Gene regulation and speciation in house mice. Genome Res 26:451–461.

Marlin BJ, Mitre M, D’Amour J A, Chao MV, Froemke RC (2015) Oxytocin enables maternal behaviour by balancing cortical inhibition. Nature 520:499–504.

McGraw LA, Davis JK, Thomas PJ, Young LJ, Thomas JW (2012) BAC-based sequencing of behaviorally-relevant genes in the prairie vole. PLoS One 7:e29345.

McGraw LA, Davis JK, Lowman JJ, ten Hallers BFH, Koriabine M, Young LJ, de Jong PJ, Rudd MK, Thomas JW (2010) Development of genomic resources for the prairie vole (Microtus ochrogaster): construction of a BAC library and vole-mouse comparative cytogenetic map. BMC Genomics 11:70.

Mitre M, Marlin BJ, Schiavo JK, Morina E, Norden SE, Hackett TA, Aoki CJ, Chao MV, Froemke RC (2016) A Distributed Network for Social Cognition Enriched for Oxytocin Receptors. J Neurosci 36:2517–2535.

Newmaster KT, Nolan ZT, Chon U, Vanselow DJ, Weit AR, Tabbaa M, Hidema S, Nishimori K, Hammock EAD, Kim Y (2020) Quantitative cellular-resolution map of the oxytocin receptor in postnatally developing mouse brains. Nature Communications 11:1885.

Olazábal DE, Young LJ (2006) Oxytocin receptors in the nucleus accumbens facilitate “spontaneous” maternal behavior in adult female prairie voles. Neuroscience 141:559–568.

Osada N, Miyagi R, Takahashi A (2017) Cis- and Trans-regulatory Effects on Gene Expression in a Natural Population of Drosophila melanogaster. Genetics 206:2139–2148.

Parmaksiz D, Kim Y (2025) Navigating Central Oxytocin Transport: Known Realms and Uncharted Territories. Neuroscientist 31:234–261.

Phelps SM, Young LJ (2003) Extraordinary diversity in vasopressin (V1a) receptor distributions among wild prairie voles (Microtus ochrogaster): patterns of variation and covariation. J Comp Neurol 466:564–576.

Rajderkar S et al. (2023) Topologically associating domain boundaries are required for normal genome function. Communications Biology 6:435.

Rangel MJ, Baldo MVC, Canteras NS (2018) Influence of the anteromedial thalamus on social defeat-associated contextual fear memory. Behavioural Brain Research 339:269–277.

Rigney N, de Vries GJ, Petrulis A, Young LJ (2022) Oxytocin, Vasopressin, and Social Behavior: From Neural Circuits to Clinical Opportunities. Endocrinology 163.

Rogers Flattery CN, Coppeto DJ, Inoue K, Rilling JK, Preuss TM, Young LJ (2022) Distribution of brain oxytocin and vasopressin V1a receptors in chimpanzees (Pan troglodytes): comparison with humans and other primate species. Brain Struct Funct 227:1907–1919.

Ross HE, Young LJ (2009) Oxytocin and the neural mechanisms regulating social cognition and affiliative behavior. Front Neuroendocrinol 30:534–547.

Ross HE, Freeman SM, Spiegel LL, Ren X, Terwilliger EF, Young LJ (2009) Variation in oxytocin receptor density in the nucleus accumbens has differential effects on affiliative behaviors in monogamous and polygamous voles. J Neurosci 29:1312–1318.

Schmidt EF, Kus L, Gong S, Heintz N (2013) BAC transgenic mice and the GENSAT database of engineered mouse strains. Cold Spring Harb Protoc 2013.

Shenoy SA, Zheng S, Liu W, Dai Y, Liu Y, Hou Z, Mori S, Tang Y, Cheng J, Duan W, Li C (2022) A novel and accurate full-length HTT mouse model for Huntington’s disease. Elife 11:e70217.

Shi X, Ng DWK, Zhang C, Comai L, Ye W, Jeffrey Chen Z (2012) Cis- and trans-regulatory divergence between progenitor species determines gene-expression novelty in Arabidopsis allopolyploids. Nature Communications 3:950.

Signor SA, Nuzhdin SV (2018) The Evolution of Gene Expression in cis and trans. Trends Genet 34:532–544.

Skuse DH, Lori A, Cubells JF, Lee I, Conneely KN, Puura K, Lehtimäki T, Binder EB, Young LJ (2014) Common polymorphism in the oxytocin receptor gene (*OXTR*) is associated with human social recognition skills. Proceedings of the National Academy of Sciences 111:1987–1992.

Smirnov A, Fishman V, Yunusova A, Korablev A, Serova I, Skryabin BV, Rozhdestvensky TS, Battulin N (2019) DNA barcoding reveals that injected transgenes are predominantly processed by homologous recombination in mouse zygote. Nucleic Acids Research 48:719–735.

Stolzenberg DS, Stevens JS, Rissman EF (2012) Experience-facilitated improvements in pup retrieval; evidence for an epigenetic effect. Horm Behav 62:128–135.

Suster ML, Abe G, Schouw A, Kawakami K (2011) Transposon-mediated BAC transgenesis in zebrafish. Nature Protocols 6:1998–2021.

Taguchi T et al. (2019) α -Synuclein BAC transgenic mice exhibit RBD-like behaviour and hyposmia: a prodromal Parkinson’s disease model. Brain 143:249–265.

Takayanagi Y, Yoshida M, Bielsky IF, Ross HE, Kawamata M, Onaka T, Yanagisawa T, Kimura T, Matzuk MM, Young LJ, Nishimori K (2005) Pervasive social deficits, but normal parturition, in oxytocin receptor-deficient mice. Proceedings of the National Academy of Sciences 102:16096–16101.

Theofanopoulou C, Andirkó A, Boeckx C, Jarvis ED (2022) Oxytocin and vasotocin receptor variation and the evolution of human prosociality. Comprehensive Psychoneuroendocrinology 11:100139.

Venkatesh B, Si-Hoe SL, Murphy D, Brenner S (1997) Transgenic rats reveal functional conservation of regulatory controls between the Fugu isotocin and rat oxytocin genes. Proc Natl Acad Sci U S A 94:12462–12466.

Walum H, Young LJ (2018a) The neural mechanisms and circuitry of the pair bond. Nat Rev Neurosci 19:643–654.

Walum H, Young LJ (2018b) The neural mechanisms and circuitry of the pair bond. Nature Reviews Neuroscience 19:643–654.

Wang Y, Song F, Zhang B, Zhang L, Xu J, Kuang D, Li D, Choudhary MNK, Li Y, Hu M, Hardison R, Wang T, Yue F (2018) The 3D Genome Browser: a web-based browser for visualizing 3D genome organization and long-range chromatin interactions. Genome Biol 19:151.

Wendt KS, Yoshida K, Itoh T, Bando M, Koch B, Schirghuber E, Tsutsumi S, Nagae G, Ishihara K, Mishiro T, Yahata K, Imamoto F, Aburatani H, Nakao M, Imamoto N, Maeshima K, Shirahige K, Peters JM (2008) Cohesin mediates transcriptional insulation by CCCTC-binding factor. Nature 451:796–801.

Wolff M, Vann SD (2019) The Cognitive Thalamus as a Gateway to Mental Representations. J Neurosci 39:3–14.

Xiao JY, Hafner A, Boettiger AN (2021) How subtle changes in 3D structure can create large changes in transcription. Elife 10:e64320.

Yang XW, Lu X-H (2008) Chapter 19 - The Bac Transgenic Approach to Study Parkinson’s Disease in Mice. In: Parkinson’s Disease (Nass R, Przedborski S, eds), pp 247-268. San Diego: Academic Press.

Young LJ, Crews D (1995) Comparative neuroendocrinology of steroid receptor gene expression and regulation: Relationship to physiology and behavior. Trends Endocrinol Metab 6:317–323.

Young LJ, Zhang QI (2021) On the Origins of Diversity in Social Behavior. The Japanese Journal of Animal Psychology doi: 10.2502/janip.71.1.4.

Zhang Q, Sano C, Masuda A, Ando R, Tanaka M, Itohara S (2016) Netrin-G1 regulates fear-like and anxiety-like behaviors in dissociable neural circuits. Scientific Reports 6:28750.

